# Mitigating CYP450-Mediated Insecticide Resistance in Malaria Vectors with Cannabis-Derived Synergists: The Potential of Cannabidiol

**DOI:** 10.64898/2026.02.10.705034

**Authors:** Maria Chalkiadaki, Linda Grigoraki, Dimitra Tsakireli, Georgia Vasalaki, Petros S. Tzimas, Mengling Chen, Latifa Remadi, Rino Ragno, Ifigeneia Akrani, Emmanouel Mikros, Rafaela Panteleri, Spyridon Vlogiannitis, Vassilios Myrianthopoulos, Ioannis K. Kostakis, Leandros A. Skaltsounis, John Vontas, Maria Halabalaki

**Affiliations:** Department of Pharmacy, National and Kapodistrian University of Athens, Panepistimiopolis Zografou, Athens, 15771, Greece; Institute of Molecular Biology and Biotechnology, Foundation for Research and Technology-Hellas, 73100 Heraklion, Greece; Department of Crop Science, Agricultural University of Athens, 11855 Athens, Greece; Rome Center for Molecular Design, Department of Drug Chemistry and Technology, Sapienza University of Rome, Piazzale Aldo Moro 5, 00185 Rome, Italy; Pharma-Informatics Unit, Athena Research and Innovation Center in Information Communication and Knowledge Technologies, Marousi, Greece

**Keywords:** cannabidiol, insecticide resistance, metabolic detoxification, insecticide synergists, *Anopheles gambiae*, CYP9K1, semisynthesis

## Abstract

Insecticide resistance in mosquitoes, largely mediated by cytochrome P450 monooxygenases (CYPs), compromises the efficacy of vector control tools. In this study, chemically-wise selected natural extracts and compounds were screened for their CYP inhibition potential. Among 37 tested plant extracts and fractions, a decarboxylated acidic fraction of industrial hemp (*Cannabis sativa* Linnaeus var. ‘Futura 75’) emerged as a promising hit, and phytochemical profiling identified cannabidiol (CBD) as its major component (IC₅₀ = 18.37 μM for CYP9K1). CBD was used as a scaffold to generate semisynthetic analogues; of which a piperazinyl analogue outperformed the natural scaffold demonstrating significantly greater potency (IC₅₀ = 2.50 μM for CYP9K1). Docking studies using homology-derived CYP9K1 models also supported a stronger binding affinity of the piperazinyl analogue relative to CBD. Toxicity assays using pyrethroid-resistant *Anopheles gambiae* Giles adults confirmed that neither CBD nor the piperazinyl analogue had intrinsic toxicity, yet the semisynthetic analogue significantly enhanced deltamethrin efficacy, showing a threefold synergistic effect. The safety profile of the cannabis compounds for non-target organisms was evaluated through human cell line cytotoxicity tests and bee toxicity assays, suggesting low non-target organism toxicity. Our study describes the identification of a plant-derived synergist lead with strong potential as an insecticide additive to combat metabolic resistance in malaria-transmitting mosquitoes.

## 1. Introduction

Cytochrome P450s are a large enzyme family with a role in diverse physiological processes, but are most well known for their ability to metabolise endogenous and exogenous compounds (Nauen et al., 2022). They act mostly through hydroxylating their substrates to less toxic and more water-soluble (and thus more readily excretable) compounds. P450s have been implicated in the adaptation of insects to plant-derived toxins, allowing them to feed on their host plants (Heidel-Fischer and Vogel, 2015; Li et al., 2007; Schuler, 2011), but critically also underlie in many cases the selection of resistance to chemical insecticides.

Insecticide resistance is a major threat to the remarkable gains achieved over the last decades in controlling insect populations that are either crop pests or disease vectors. In the case of *Anopheles gambiae* Giles mosquitoes, the main vectors of malaria (a disease that caused over 597,000 deaths in 2023; World Health Organization, 2024) resistance to most commonly used insecticides is now widespread. Particularly concerning is the selection of resistance to pyrethroid insecticides, that are used in all insecticide treated bed nets (ITNs), with limited alternative options available. Pyrethroid resistance in *An. gambiae* is known to be linked with the over-expression of P450s, particularly members of the CYP6 and CYP9 clades (Vontas et al., 2020). One of the best characterised pyrethroid-metabolising P450s is CYP9K1, which is over-expressed in several resistant populations (Hearn et al., 2022; Sandeu et al., 2020; Vontas et al., 2018) and in *vitro* (Vontas et al., 2018) and *in vivo* (unpublished) data have verified its ability to metabolise pyrethroids and confer resistance.

To alleviate P450 based insecticide resistance, current interventions rely on mixtures of insecticides with synergists, i.e. non-toxic compounds able to inhibit detoxification enzymes and thereby restore insecticidal efficacy (World Health Organization, 2022). Piperonyl butoxide (PBO), a broad-spectrum CYP450 inhibitor, is widely used in combination with pyrethroids in ITNs to counteract metabolic resistance in mosquito populations. Since the World Health Organization (WHO) endorsed pyrethroid–PBO bed nets in 2017 (World Health Organization, 2017), their use has improved vector control in regions with high pyrethroid resistance (Bayili et al., 2019; Churcher et al., 2016; Corbel et al., 2010; Gleave et al., 2021; Koudou et al., 2011; Menze et al., 2020; N’Guessan et al., 2010; Oumbouke et al., 2019; Pennetier et al., 2013; Syme et al., 2022; Toe et al., 2018; Tungu et al., 2010, 2021). Nonetheless, reports of reduced PBO synergy in *An. gambiae* field populations (Oruni et al., 2024; Pelloquin et al., 2025) raise concerns that extensive use of this compound could select for insensitivity and ultimately lead to a loss of PBO’s operational efficacy. Thus, expanding the arsenal of synergists is of paramount importance to provide alternative options for effective resistance management.

Plants are natural chemical factories, producing a vast array of secondary metabolites that serve as innate defenses against herbivores (Schuler, 2011). While over 2,000 plant species are known to possess insecticidal properties, only a small subset has been thoroughly studied (Edwin et al., 2016). Beyond their direct toxicity, several plant-derived extracts have been shown to act as insecticide synergists by inhibiting detoxification enzymes (Bullangpoti et al., 2011; Francis et al., 2025; Kotewong et al., 2014; Mohan et al., 2006; Pethuan et al., 2012). Mediterranean taxa, shaped by recurring drought and high irradiance, frequently exhibit chemically diverse metabolomes that support defense and stress tolerance (Qaderi et al., 2023), providing an unexplored pool of compounds for exploring plant-derived synergists targeting metabolic resistance.

This study investigates the potential of thirty-seven extracts/fractions from twenty-one plant species from Mediterranean/arid environments with chemically diverse defensive metabolomes, and industrially scalable sources to inhibit cytochrome P450 enzymes, that are major insecticide metabolisers. Screening of the plant extracts was followed by phytochemical analysis and *in vivo* toxicity and pyrethroid synergism testing. Semisynthetic optimisation produced a more potent derivative of the main active compound as shown by *in vitro*, *in vivo* and *in silico* assays. The toxicity profile of the compounds to non-target organisms was also addressed. By exploring novel CYP450-modulating synergists from natural sources, this research contributes new tools for mitigating insecticide resistance and enhancing the efficacy of mosquito control strategies.

## 2. Materials and methods

### 2.1. Materials and chemicals

The fluorescent substrate and product 7-ethoxy-coumarin (7-EC) and 7-hydroxy-coumarin (7-HC), as well as anhydrous sodium phosphate dibasic (Na_2_HPO4), anhydrous sodium phosphate monobasic (NaH_2_PO4), Glutathione reductase from baker’s yeast (*S. cerevisiae*), tris(hydroxy-methyl)aminomethane (Trizma© base), quercetin and the high-performance liquid chromatography (HPLC)-grade chemicals β-Νicotinamide adenine dinucleotide 2’-phosphate reduced tetrasodium salt hydrate (NADPH) (≥93%), L- Glutathione oxidized (≥98%), were purchased from Sigma-Aldrich, Merck KGaA, Darmstadt, Germany.

Water (H_2_O) was obtained from a Millipore Direct-Q 3 UV purification system (Merck KGaA, Darmstadt, Germany), acetonitrile (ACN) liquid chromatography-mass spectrometry (LC-MS) grade was purchased from Supelco, Merck KGaA, Darmstadt, Germany. ACN HPLC grade (≥99.9%), ethanol (EtOH) absolute HPLC grade (≥99.8%) and formic acid (≥99.0%) Optima™ LC-MS grade were purchased from Fisher Chemical, Fisher Scientific, Waltham, MA, USA. Hydrochloric acid (HCl) (fuming, ≥37%) was purchased from Honeywell Fluka™ Charlotte, NC, USA. Formic acid (99-100% a.r.) was purchased from AnalytiChem Belgium NV, Zedelgem, Belgium. The 96- well plates were purchased from Greiner Bio-One (Frickenhausen, Germany).

For the semisynthetic reactions, 2-butanone (≥99.0%), diethyl ether (EMPARTA^®^, 99.5%) piperazine (ReagentPlus^®^, 99%), 3,3΄-diamino-N-methyldipropylamine (96%) were purchased from Sigma-Aldrich, Merck KGaA, Darmstadt, Germany. Caesium carbonate (Cs_2_CO_3_, 99.0%), 1,6-diaminohexane (98.0%), 1,4-diaminobutane (98.0%), and ethyl bromoacetate (95%) were purchased from Fluorochem Ltd. (Hadfield, UK). Dichloromethane (≥99.9%), n-hexane (≥95%), and methanol, (MeOH, ≥99.9%), were purchased from Carlo Erba Reagents, Milan, Italy. Sodium chloride (NaCl) was purchased from Fisher Chemical, Fisher Scientific, Waltham, MA, USA. Triethylamine (Et_3_N) was obtained from Penta Chemicals Unlimited, Prague, Czech Republic. Ethylenediamine and 1,10-diaminodecane were purchased from Honeywell Fluka™ (Honeywell, Charlotte, NC, USA).

For the cytotoxicity assays on mammalian cells, the following reagents were used: Dulbecco’s Modified Eagle Medium (DMEM, low glucose), penicillin-streptomycin (10,000 U/mL), and fetal bovine serum were purchased from Gibco™, Fisher Scientific (Madrid, Spain). Eagle’s Minimum Essential Medium (EMEM), L -glutamine, non-essential amino acids and sodium pyruvate were obtained from Merck (Merck KGaA, Darmstadt, Germany). 3-(4,5-dimethylthiazol-2-yl)-2,5-diphenyltetrazolium bromide (MTT) and dimethyl sulfoxide (DMSO) were purchased from Sigma Aldrich (Merck KGaA, Darmstadt, Germany).

### 2.2. Selection and source of plant extracts

Extracts selected for the initial *in vitro* screening sourced from the in-house natural products repository of the Laboratory of Valorization of Bioactive Natural Products, NKUA. From the repository’s collection of over 2,000 extracts derived from the Greek, Mediterranean and global biodiversity, a subset was selected based on two primary criteria: (1) documented insecticidal activity in prior studies for the source species, and (2) representation of major, structurally diverse phytochemical classes as determined by preliminary UHPLC-HRMS metabolite profiling (data not shown). The complete list of selected source species and their corresponding extract types (e.g., crude methanol or ethyl acetate extracts from ultrasound-assisted extraction (UAE), fractions from liquid-liquid partitioning, XAD-7 resin fractions, and enriched extracts from supercritical fluid extraction (SFE)) is provided in Supplementary Table S1.

Hemp extracts of *C. sativa* ‘Futura 75’ (SFE or EtOH-UAE) were fractionated by a two-stage pH-controlled liquid–liquid extraction (LLE) to selectively enrich cannabinoids. The total extracts were dissolved in n-hexane/ethyl acetate (80:20, *v/v*) and partitioned against ethanol/water (50:50, *v/v*). Alkalinisation of the aqueous phase to pH 10 with aqueous ammonia converted the naturally abundant acidic cannabinoids (e.g., CBDA) into water-soluble salts, which partitioned into the aqueous layer, while non-ionisable lipophilic components (including lipids and waxes) remained in the organic phase (neutral fraction). The aqueous phase was then contacted with fresh organic solvent and acidified to pH ≈ 1 with hydrochloric acid, converting the salts back to non-ionised acidic cannabinoids that were back-extracted into the organic layer (acidic fraction). This fraction, reflecting the naturally abundant form of cannabinoids, was subsequently thermally decarboxylated (150 °C, 2 h) to yield a fraction enriched in neutral cannabinoids, mostly CBD (acidic fraction decarboxylated). All organic fractions were washed with distilled Η_2_Ο and were concentrated to dryness.

### 2.3. Functional expression of major cytochrome P450 insecticide metabolisers

The expression of *Bt*CYP6CM1 and *Αg*CYP9K1 was carried out following previously published protocols (Karunker et al., 2009; Vontas et al., 2018). Cytochrome P450 gene sequences were synthesised and cloned into the pCW-OmpA2 expression vector, while *Anopheles gambiae* cytochrome P450 reductase (*Ag*CPR) was cloned into the pACYC vector. Recombinant plasmids were co-transformed into *E. coli* BL21STAR cells.

Transformed cells were grown in terrific broth with appropriate antibiotic selection and induced with δ-aminolevulinic acid (Melford Laboratories Ltd., Ipswich, UK) and isopropyl β-D-1-thiogalactopyranoside (IPTG; Sigma-Aldrich, Merck KGaA, Darmstadt, Germany). Cells were harvested, converted into spheroplasts, and membrane fractions were prepared by differential centrifugation. Membrane preparations were stored at −80 °C and analysed for total protein content, functional P450 expression by CO-difference spectroscopy, and CPR activity via cytochrome c reduction. *An. gambiae* cytochrome b5 was produced as previously described (Stevenson et al., 2011).

### 2.4. *In vitro* inhibition activity of CYP6CM1 and CYP9K1 using fluorescent model substrate

Plant extracts with reported insecticidal properties, as well as those containing known major compounds representing distinct chemical classes, were selected for screening against cytochrome P450 enzymes CYP6CM1 from *Bemisia tabaci* (Gennadius) and CYP9K1 from *Anopheles gambiae. Bt*CYP6CM1 was used for the initial screening, given the greater availability of this recombinant enzyme, which also shows abroad metabolic capacity. Extracts identified as active were then evaluated for inhibition of *Ag*CYP9K1.

Enzyme activity reactions were conducted in black, flat-bottom 96-well plates using bacterial membrane preparations. Each reaction (final volume 120 μL) contained 10 pmol of recombinant CYP6CM1vQ or *Ag*CYP9K1 and 100 pmol of *Anopheles gambiae* cytochrome b₅ in 0.1 M sodium phosphate buffer (pH 7.2). Cytochrome P450 activity was monitored using the fluorogenic substrate 7-ethoxycoumarin (7-EC), following a previously described protocol with minor modifications (Karunker et al., 2009).

Test extracts were dissolved in ethanol and added to the reactions at a final concentration of 8.33 μg/mL with a final ethanol concentration of 8 % (*v/v*), which was kept constant across all reactions. The substrate 7-EC was added to a final concentration of 0.83 mM. Plates were pre-warmed for 10 minutes at 30 °C and reactions were initiated by the addition of NADPH to a final concentration of 1.25 mM. Following incubation for 30 min at 30 °C, background fluorescence resulting from residual NADPH was eliminated by the addition of oxidised glutathione (30 mM) and glutathione reductase (0.125 U per well). After a 10-min incubation at room temperature, reactions were terminated by adding 140 mM of a 50:50 (*v/v*) acetonitrile/Tris-HCl buffer (pH 8.5).

The formation of the fluorescent O-dealkylated product, 7-hydroxycoumarin (umbelliferone), was quantified by measuring fluorescence at 390 nm excitation and 465 nm emission using a Tecan Infinite 200 PRO plate reader (Tecan, Männedorf, Switzerland).

A standard curve of 7-hydroxycoumarin was generated by serial dilution of a stock solution in ethanol and measured under identical conditions. This standard curve was used to convert fluorescence units obtained from the 7-ethoxycoumarin O-deethylation assay into absolute amounts of product formed, enabling quantitative comparison of enzymatic activity between treatments.

Negative controls lacking NADPH were included to account for non-enzymatic background fluorescence, while reactions without inhibitors served as activity controls. Solvent controls containing 8% (*v/v*) ethanol were included to correct for solvent effects. Quercetin was used as a positive control at a final concentration of 26 μM. All assays were performed in triplicate.

IC₅₀ values were determined from dose–response curves generated using a range of concentrations for CBD and its more potent derivative, CBD analogue 3. Nonlinear regression analysis was performed using GraphPad Prism version 8.0.1 (GraphPad Software Inc., San Diego, CA, USA). Serial dilutions of each compound were prepared in ethanol (8% *v/v*, constant across all treatments), over a concentration range of 0.01–200 µM for Analogue 3 and 1–500 µM for CBD.

### 2.5. Isolation of cannabidiol and cannabidiolic acid

Cannabidiolic acid (CBDA) was isolated from a hemp (*Cannabis sativa* L., variety ‘Futura 75’) extract, obtained via supercritical fluid extraction (SFE) with CO_2_. Purification was achieved using an established centrifugal partition chromatographic (CPC) method (Popp et al., 2019). Cannabidiol (CBD) was subsequently isolated after decarboxylation of CBDA, following a modified purification protocol, as described by Tzimas et al., 2021.

Structural confirmation of both compounds was achieved through liquid chromatography–high-resolution mass spectrometry (LC-HRMS) and one- and two-dimensional nuclear magnetic resonance (NMR) spectroscopy, compared with literature data (Berman et al., 2018; Choi et al., 2004; Citti et al., 2019). Compound purity was determined to be >99% by ultra-high-performance liquid chromatography with photodiode array detection (UPLC-PDA) on an Acquity UPLC-PDA system (Waters, Milford, MA, USA). CBD and CBDA analytical standards were obtained in-house. Stock solutions of CBD and CBDA (4 mg/mL) were prepared by dissolving accurately weighed amounts in ACN. For quantitative analysis, mixed standard working solutions were prepared in the same solvent. All standard solutions were stored at -20 °C.

### 2.6. Differential scanning fluorimetry (DSF)

Experiments were performed in a BioRad CFX-Connect real time PCR machine. The DSF protocol (Wu et al., 2023) involved an initial 3 min sample incubation at 25 °C and subsequent heating at a continuous rate of 0.5 °C per minute repeated for 65 cycles. The protein was diluted at a concentration of 2 μM in a buffer comprising 10 mM HEPES at pH 7.4 and 150 mM NaCl, while for monitoring protein unfolding the ProteOrange stain reagent (Lumiprobe code 10210) was used at 10x concentration. The ligand was diluted in the protein buffer from a 10 mM stock solution in 100% DMSO and was assayed at a concentration of 250 μM. Analysis and calculation of Δ*T*_m_ was performed by BioRad Maestro software and the fitting tools of the DSF-World application.

### 2.7. Phytochemical analysis

#### 2.7.1. Ultra high-performance liquid chromatography–photodiode array (UPLC-PDA) quantification and purity analysis of cannabinoids

Quantitative and purity analysis of the extract with the strongest inhibitory activity was conducted to determine its exact composition and confirm that its components are responsible for CYP450 inhibition. For this purpose, an Acquity UPLC system equipped with a photo diode array (PDA) detector (Waters, Milford, MA, USA) was utilised. Waters Empower 3 software was used for instrument control, data acquisition and processing.

Chromatographic separation was performed on a C_18_-PFP core-shell column (SpeedCore, 100 mm × 2.1 mm, 2.6 μm; Fortis Technologies, Neston, Cheshire, UK). The mobile phase consisted of (A) H_2_O and (B) ACN, both containing 0.1% formic acid and the analysis was conducted as described by Tzimas et al., 2024. Briefly, the flow rate was set at 0.4 mL/min. The elution program was as follows: 60% B for 0.5 minutes, a linear increase to 68% B over 2.4 minutes, isocratic at 68% B for 2.6 minutes, a linear increase to 90% B over 2.3 minutes, and isocratic at 90% B for 1.2 minutes. The initial composition was restored within 0.1 minutes and maintained for 1.9 minutes for column re-equilibration, resulting in a total run time of 11 minutes. The column compartment and autosampler were kept at 30 °C and 10 °C, respectively. The injection volume was 3 μL.

PDA spectra were recorded in the range 200–500 nm, and chromatograms for quantitative analysis were obtained at selected wavelengths (210 and 225 nm) based on the absorption maxima (*λ*_max_) of each analyte. External standard calibration curves were constructed for the quantification of cannabinoids. Compound identification was based on the retention time and the UV-Vis spectrum in comparison to standards.

#### 2.7.2. Ultra high-performance liquid chromatography–high-resolution tandem mass spectrometry (UHPLC-HRMS/MS) qualitative profiling and compound identification

Qualitative determination of the most potent extract was conducted using ultra high-performance liquid chromatography–high-resolution mass spectrometry (UHPLC-HRMS) and tandem mass spectrometry (HRMS/MS) experiments to annotate the major constituents and support identification of the active compound. The analysis was performed on a Vanquish UHPLC system coupled to an Orbitrap Exploris 120 mass spectrometer (both Thermo Fisher Scientific, Waltham, MA, USA). Chromatographic separation was achieved using a CORTECS UPLC C18 column (150 mm x 2.1 mm, 1.6 μm; Waters, Milford, MA, USA), maintained at 40 ^O^C.

The mobile phase consisted of (A) H_2_O containing 0.1% formic acid and (B) ACN. The elution gradient was as follows: 15% B for 0.5 minutes, increased to 65% B over the next 1 minute, then linearly increased to 66% in the next 16.5 minutes, reached 100% in the next 7 minutes and was maintained isocratically for 2 minutes. Finally, the system returned to the initial conditions for re-equilibration. The total acquisition time was 30 minutes, with a flow rate of 300 μL/min. The injection volume was 5 μL and the autosampler temperature was set at 7 °C. Samples were prepared at a concentration of 300 μg/mL using a MeOH/ H_2_O (90:10, *v/v*) solution as the dilution solvent.

Mass spectra were acquired in positive and negative ionisation modes using a heated electrospray ionisation (HESI) source. Source parameters were as follows: vaporiser temperature, 350 °C; ion transfer tube temperature, 350 °C; spray voltage, 3.8 kV (positive mode) and 3.6 kV (negative mode); sheath gas, 45 arbitrary units; auxiliary gas, 20 arbitrary units; sweep gas, 0 arbitrary units.

The HRMS data were acquired in full scan mode over an *m/z* of 113-800, with a resolving power of 60,000 (full width at half maximum, FWHM, at *m/z* 200). HRMS/MS experiments were conducted at a resolving power of 30,000 (FWHM at *m/z* 200) using a top-3 data-dependent acquisition (DDA) mode with normalised collision energy steps of 30%, 50% and 150%. Data acquisition was performed using Xcalibur 4.6 software and data processing was carried out using Freestyle 1.8 software (Thermo Fisher Scientific, Waltham, MA, USA).

Cannabinoid identification was performed by examining the base peak (BP) chromatograms and extracted ion chromatograms (EICs) to obtain the corresponding full-scan spectra. The Freestyle software calculator tool was used for the proposed elemental composition of each *m/z* value, with a mass error below 5 ppm, supported by isotopic patterns and ring double-bond equivalent (RDBeq) values. Cannabinoids were identified by comparison with authentic reference standards based on retention time, exact mass, and HRMS/MS spectra acquired under identical analytical conditions.

### 2.8. Semisynthesis of target compounds

#### 2.8.1. Ethyl 2-(((1’R,2’R)-6-hydroxy-5’-methyl-4-pentyl-2’-(prop-1-en-2-yl)-1’,2’,3’,4’-tetrahydro-[1,1’-biphenyl]-2-yl)oxy)acetate (Analogue 1)

A solution of CBD (1.57 g, 1 eq, 4.99 mmol) in butanone (50 mL) was stirred at room temperature (RT) for 15 min under an argon (Ar) atmosphere. Then, Cs₂CO₃ (2.44 g, 1.5 eq, 7.49 mmol) was added and the resulting mixture was stirred at 90°C for 1 h under Ar. Subsequently, ethyl bromoacetate (1.08 g, 1.3 eq, 6.49 mmol) was added and the reaction was allowed to proceed at 90°C under continuous stirring for 16 h. Upon completion, the reaction mixture was concentrated under reduced pressure. The resulting solid residue was dissolved in a mixture of diethyl ether: n-hexane (2:1 *v/v*, 80 mL) and washed successively with H₂O (3 × 50 mL) and a saturated NaCl solution (40 mL). The organic phase was collected, dried over anhydrous Na₂SO₄, filtered through a fluted filter and concentrated under vacuum to dryness. The crude product was purified by column chromatography (aluminium oxide 90 active – Brockman III) using diethyl ether: n-hexane (1:49 *v/v*) as the mobile phase, affording pure CBD Analogue 1. The isolated product was obtained as a solid (1 g, yield ∼58%).

A solution of CBD (1.57 g, 4.99 mmol) in butanone (50 mL) was stirred at room temperature (RT) for 15 min under an argon atmosphere. Cs₂CO₃ (2.44 g, 7.49 mmol) was then added, and the resulting mixture was stirred at 90°C for 1 h under argon. Ethyl bromoacetate (1.08 g, 6.49 mmol) was subsequently added, and the reaction mixture was stirred at 90°C for 16 h. Upon completion of the reaction, the mixture was concentrated under reduced pressure, dissolved in a mixture of diethyl ether and n-hexane (2:1, 80 mL) and washed successively with water (3 × 50 mL) and saturated aqueous NaCl solution (40 mL). The organic layer was collected, dried over anhydrous Na₂SO₄, filtered, and concentrated to dryness. The crude product was purified by column chromatography (aluminium oxide, Brockmann III) using diethyl ether/n-hexane (1:49, v/v) as the eluent, to afford 1 g of Analogue 1 (58%).

^1^H NMR (600 MHz, MeOD) *δ* 6.25 (s, 1H, H-3΄), 6.12 (s, 1H, H-5΄), 5.23 (br. s, 1H, H-2), 4.54 (d, J_11a-11b_ = 15.7 Hz, 1H, H-11_a_), 4.49 – 4.40 (m, 3H, H-9_cis_/ H-9_trans_/ H-11_b_), 4.23 (q, J_13-14_ = 7.1 Hz, 2H, H-13), 4.00 (m, 1H, H-1), 2.99 (m, 1H, H-6), 2.44 (t, J_a΄-b΄_ = 7.5 Hz, 2H, H-a΄), 2.22 (m, 1H, H-4), 1.99 (m, 1H, H-4), 1.74 (m, 2H, H-5), 1.66 (s, 3H, H-7), 1.62 (s, 3H, H-10), 1.56 (p, J_b΄-a΄,c΄_ = 7.7 Hz, 2H, H-b΄), 1.33 (m, 2H, H-d΄), 1.28 (t, J_14-13_ = 7.1 Hz, 3H, H-14), 0.90 (t, J_e΄-d΄_ = 7.2 Hz, 3H, H-e΄). ^13^C NMR (151 MHz, MeOD) *δ* 171.09 (C-12), 158.89 (C-2΄), 157.57 (C-6΄), 150.42 (C-8), 142.96 (C-4΄), 134.19 (C-3), 127.16 (C-2), 118.61 (C-1΄), 110.65 (C-3΄), 110.55 (C-9), 105.59(C-5΄), 67.40 (C-11), 62.08 (C-13), 46.30 (C-6), 37.56 (C-1), 36.81 (C-a΄), 32.61 (C-c΄), 32.05 (C-b΄), 31.69 (C-4), 30.81 (C-5), 23.70 (C-7), 23.59 (C-d΄), 19.50 (C-10), 14.55 (C-14), 14.40 (C-e΄). HRMS (ESI-) *m/z* 399.2557 (C_25_H_36_O_4_). 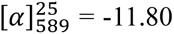 (2.0 g/ 100 mL, MeOH).

#### 2.8.2. N-(4-aminobutyl)-2-(((1’R,2’R)-6-hydroxy-5’-methyl-4-pentyl-2’-(prop-1-en-2-yl)-1’,2’,3’,4’-tetrahydro-[1,1’-biphenyl]-2-yl)oxy)acetamide (Analogue 2)

A mixture of Analogue **1** (200 mg, 0.49 mmol) and 1,4-diaminobutane (442 mg, 7.35 mmol) in ethanol (5 mL) was stirred under an argon atmosphere at 93 °C for 16 h. Upon completion of the reaction, the mixture was concentrated under reduced pressure, dissolved in dichloromethane (40 mL) and washed with water (2 × 25 mL). The organic phase was collected, dried over anhydrous Na₂SO₄, filtered, and concentrated to dryness. The residue was purified by flash chromatography (silica gel) using MeOH/Et₃N/DCM (1:1:48, v/v) as the eluent to afford 179 mg (91%) of Analogue **2** (78%).

^1^H NMR (600 MHz, MeOD) *δ* 6.30 (s, 1H, H-3΄), 6.20 (s, 1H, Η-5΄), 5.36 (br. s, 1H, H-2), 4.45 – 4.34 (m, 4H, H-9_cis_/ H-9_trans_/ H-11_a_/ H-11_b_), 4.01 (m, 1H, H-1), 3.32 (m, 2H, H-13), 2.83 (m, 1H, H-6), 2.70 (t, J_16-15_ = 7.1 Hz, 2H, H-16), 2.46 (t, J_a΄-b΄_ = 7.7 Hz, 2H, H-a΄), 2.18 (m, 1H, H-4_a_), 2.07 (m, 1H, H-4_b_), 1.79 (m, 2H, H-5), 1.72 (s, 3H, H-7), 1.62 (s, 3H, H-10), 1.59 – 1.48 (m, 6H, H-b΄/ Η-14/ H-15), 1.37 – 1.28 (m, 4H, H-d΄/ H-c΄), 0.90 (t, J = 7.0 Hz, 3H, H-e΄). ^13^C NMR (151 MHz, MeOD) *δ* 171.74 (C-12), 158.36 (C-2΄), 157.57 (C-6΄), 150.33 (C-8), 143.45 (C-4΄), 132.37 (C-3), 128.74 (C-2), 118.05 (C-1΄), 110.78 (C-9/ C-3΄), 105.53 (C-5΄), 68.66 (C-11), 53.61 (C-13), 41.21 (C-6), 39.77 (C-16), 37.84 (C-1), 36.81 (C-a΄), 32.62 (C-c΄), 32.07 (C-b΄), 31.78 (C-4), 30.65 (C-5), 28.38 (C-14), 28.00 (C-15), 23.73 (C-7), 23.59 (C-d΄), 19.67 (C-10), 14.40 (C-e΄). HRMS (ESI-) *m/z* 441.3070 (C_27_H_42_N_2_O_3_). 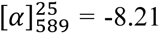 (6.2 g/ 100 mL, MeOH).

#### 2.8.3. 2-(((1’R,2’R)-6-hydroxy-5’-methyl-4-pentyl-2’-(prop-1-en-2-yl)-1’,2’,3’,4’-tetrahydro-[1,1’-biphenyl]-2-yl)oxy)-1-(piperazin-1-yl)ethan-1-one (Analogue 3)

Analogue **3 (**63%) was prepared using an analogous procedure to that of Analogue **2**, using piperazine as starting material.

^1^H NMR (600 MHz, MeOD) *δ* 6.27 (s, 1H, H-3΄), 6.24 (s, 1H, Η-5΄), 5.23 (br. s, 1H, H-2), 4.67 (d, J_11a-11b_ = 13.1 Hz, 1H,H-11_a_), 4.55 (d, J_11b-11a_ = 13.1 Hz, 1H, H-11_b_), 4.44 (dd, J_9trans-9cis_ = 0 Hz, 1H, H-9_trans_), 4.42 (dd, J_9cis-9trans_ = 0 Hz, 1H, H-9_cis_), 3.98 (m, 1H, H-1), 3.64 (m, 2H, H-13), 3.57 (m, 1H, H-14), 2.93 (m, 1H, H-6), 2.85 (m, 4H, H-15/ H-16), 2.45 (t, J_a΄-b΄_ = 7.5 Hz, 2H, H-a΄), 2.19 (m, 1H, H-4_a_), 1.99 (m, 1H, H-4_b_), 1.74 (m, 2H, H-5), 1.66 (s, 3H, H-7), 1.61 (s, 3H, H-10), 1.56 (p, J_b΄-a΄, c΄_ = 7.7 Hz, 2H, H-b΄), 1.36-1.27 (m, 4H, H-c΄/ H-d΄), 0.90 (t, Je΄-d΄ = 7.1 Hz, 3H, H-e΄). ^13^C NMR (151 MHz, MeOD) *δ* 169.46 (C-12), 159.06 (C-2΄), 157.50 (C-6΄), 150.33 (C-8), 143.16 (C-4΄), 132.78 (C-3), 127.28 (C-2), 118.31 (C-1΄), 110.61 (C-9/ C-3΄), 105.72 (C-5΄), 69.64 (C-11), 47.09 (C-13), 46.64 (C-15), 46.58 (C-6), 46.07 (C-16), 43.42 (C-14), 37.65 (C-1), 36.83 (C-a΄), 32.63 (C-c΄), 32.05 (C-b΄), 31.76 (C-4), 30.79 (C-5), 23.84 (C-7), 23.59 (C-d΄), 19.58 (C-10), 14.42 (C-e΄) HRMS (ESI-) *m/z* 439.2958 (C_24_H_40_N_2_O_3_). 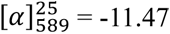 (1.5 g/ 100 mL, MeOH).

#### 2.8.4. N-(8-aminooctyl)-2-(((1’R,2’R)-6-hydroxy-5’-methyl-4-pentyl-2’-(prop-1-en-2-yl)-1’,2’,3’,4’-tetrahydro-[1,1’-biphenyl]-2-yl)oxy)acetamide (Analogue 4)

Analogue 4 (78%) was prepared using an analogous procedure to that of Analogue 2, using 1,8-diaminooctane as starting material.

^1^H NMR (600 MHz, MeOD) *δ* 6.30 (s, 1H, H-3΄), 6.19 (s, 1H, H-5΄), 5.35 (br. s, 1H, H-2), 4.45 – 4.34 (m, 4H, H-9_cis_/ H-9_trans_/ H-11_a_/ H-11_b_), 4.02 (m, 1H, H-1), 3.30 (m, 2H, H-13), 2.84 (m, 1H, H-6), 2.66 (t, J_20-19_ = 7.2 Hz, 2H, H-20), 2.45 (t, J_a΄-b΄_ = 7.6 Hz, 2H, H-a΄), 2.19 (m, 1H, H-4_a_), 2.07 (m, 1H, H-4_b_), 1.79 (m, 2H, H-5), 1.71 (s, 3H, H-7), 1.62 (s, 3H, H-10), 1.57 – 1.49 (m, 6H, Η-b΄/ H-14/ H-19), 1.37 – 1.28 (m, 12H, H-c΄/ H-d΄/ Η-15/ Η-16/ Η-17/ H-18), 0.90 (t, J_e΄-d΄_ = 7.1 Hz, 3H, Η-e΄) ^13^C NMR (151 MHz, MeOD) *δ* 171.43 (C-12), 158.11 (C-2΄), 157.55(C-6΄), 150.31 (C-8), 143.38 (C-4΄), 132.77 (C-3), 128.79 (C-2), 117.89 (C-1΄), 110.75 (C-9/ C-3΄), 105.29 (C-5΄), 68.50 (C-11), 46.97 (C-6), 42.24 (C-20), 40.22 (C-13), 37.82 (C-1), 36.85 (C-a΄), 32.94 (C-d΄), 32.64 (C-19), 32.06 (C-b΄), 31.76 (C-4), 30.70 (C-15/ C-18), 30.46 (C-5), 30.34 (C-14), 27.88 (C-16/ C-17), 23.72 (C-7), 23.59 (C-c΄), 19.66 (C-10), 14.43 (C-e΄). HRMS (ESI-) *m/z* 497.3752 (C_31_H_50_N_2_O_3_). 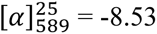 (4.8 g/ 100 mL, MeOH).

### 2.9. *An. gambiae* adult bioassays

#### 2.9.1. Mosquito strains and rearing

Mosquitoes were maintained at the following conditions: temperature of 26 ± 2 °C and 80 ± 10% relative humidity, with a 12 h:12 h light–dark photoperiod including a 1 h simulated dawn and dusk. Larvae were provided with finely ground fish food (Tetramin Tropical Flakes; Tetra, Blacksburg, VA, USA), while adults were maintained on a 10% sucrose solution ad libitum. The pyrethroid resistant VK7 strain was provided by iiDiagnostics Limited (“iiDiagnostics”), a subsidiary of the Liverpool School of Tropical Medicine and is subjected to selection with 0.05% deltamethrin (WHO test paper) at a 2h treatment every 2∼3 generations.

#### 2.9.2. Topical application bioassay

Deltamethrin (99.40%) was purchased from Dr. Ehrenstorfer (LGC Labo, Augsburg, Germany) and dissolved in acetone, while the test compounds were dissolved in ethanol. Female mosquitoes (3–5 days old) were randomly selected, briefly anesthetised on ice, and transferred to Petri dishes. A volume of 0.2 µL of each insecticide, test compound or combinatorial mixture was topically applied to the pronotum using a PB600-1 repeating dispenser (Hamilton Company, Reno, USA). Both ethanol and acetone solvents were tested for mortality in isolation showing less than 10% mortality, as well as in a 1:1 mixture that was used as control for combinatorial mixtures. Each dose was tested in at least three independent replicates, with a minimum of 10 individuals per replicate. Mortality was recorded 24 h post-treatment. Data were analysed using GraphPad Prism version 8.2.1 (GraphPad Software Inc., San Diego, CA, USA).

### 2.10. *In silico* calculations

#### 2.10.1. Potential inhibitors identification workflow

Structurally characterised homologues of CYP9K1 were identified using the online platform HHpred (MPI Bioinformatics Toolkit), which employs hidden Markov models (HMMs) for homology detection (Zimmermann et al., 2018). A compound library for docking calculations was prepared in smiles format. Potential inhibitors of CYP3A4 were predicted using three ADME prediction tools, namely SwissADME (Daina et al., 2017), ADMET Predictor (Simulation Plus Inc.), and ADMETlab 2.0 (Xiong et al., 2021). Consensus selection was performed by retaining only the compounds predicted as inhibitors by all 3 programs using R programing language to filter the results. Diversity screening was conducted using the Canvas Similarity and Clustering tool within Schrödinger maestro suite (Schrödinger Release 2021-2: Maestro, Schrödinger, LLC, New York, NY, 2021) with the default parameters selection of representative compounds nearest the centroid of each cluster.

#### 2.10.2. CBD binding prediction

Considering the high flexibility of the CYP450 active site to accommodate a wide variety of substrates, we adopted a consensus docking strategy consisting of consecutive filtering steps as following: a) Generation of adequate 3D protein structure models (because of the lack of experimental crystal structures). b) Consensus molecular docking using 15 different algorithms. This approach is based on the principle that when multiple independent docking algorithms converge on similar binding poses, the probability of identifying a reliable binding mode increase. c) Selection based on binding free energy calculations ΔG_bind_ among representative structures of the most populated clusters. d) Induced Fit Docking based on the most populated cluster of the previous step. e) Final selection based on binding free energy calculations ΔG_bind_. The details for each one of the above-mentioned steps are the following.

##### 2.10.2.1. CYP9K1 models preparation

Two structural models of CYP9K1 were obtained from the AlphaFold (Jumper et al., 2021) and MODELLER (modbase) (Pieper et al., 2014) databases and prepared for docking studies. The missing heme cofactor was extracted from the human CYP3A4 structure. This choice was guided by a BLAST (Altschul et al., 1990) search using the CYP9K1 sequence. Among the available CYP3A4 crystal structures, only seven were co-crystallized with an inhibitor (6BD7, 6UNL, 7KVM, 6BDH, 6UNM, 9COY, 6UNI, 7KVI, 6UNJ, 7KVK).Each experimental structure was aligned separately to the two predicted models, and the root-mean-square deviation (RMSD) of the binding site residues (defined by all common residues within 5 Å from all the inhibitors and heme structures) was calculated to assess structural similarity (Supplementary Table S2). Based on the RMSD values, PDB ID 6UNI was selected as the closest match for the AlphaFold model, and PDB ID 6UNL was chosen for the MODELLER model. These structures served as templates for accurate placement of the heme group, which was transferred into each CYP9K1 model via structural alignment. Hydrogens were added using UCSF Chimera (Pettersen et al., 2004), and the resulting complexes were energy-minimised for 100 iterations to relieve any steric clashes and optimise geometry.

##### 2.10.2.2. Consensus docking calculations

Considering the high flexibility of the CYP9K1 active site (as reflected in the RMSD values in Table S2) and the inherent uncertainties in the AF and modbase structural models, a consensus docking strategy was adopted. This approach is based on the principle that when multiple independent docking algorithms converge on similar binding poses, the probability of identifying a reliable binding mode increase. To implement this methodology, the online platform 3D-QSAR.com (Ragno, 2019) which provides access to several docking algorithms through its integrated Py-Docking tool was utilized. Py-Docking includes 15 docking algorithms (Smina, PLANTS, PSOVina, VinaXB, VinaSH, mPSOVina, qVina2, qVinaW, iGEMDOCK, GLAMDOCK, idock, LeDock, BetaDock, VinaLast, and iMolSDock), available to the platform developers. While standard users have access only to Smina and PLANTS (with all associated scoring functions), the full suite of methods was used in this study. The docking search space was defined by positioning a methanol probe at the center of the active site of both the AF and modbase models. CBD was docked using each program, generating 128 poses. Together with the corresponding energy-minimized structures, 256 poses were collected for consensus analysis.

##### 2.10.2.3. Clustering of ligand poses

Consensus analysis was carried out by clustering of ligand poses. For that purpose, Clustering Based on Volume Overlap tool in Maestro (Schrödinger Release 2021-2: Maestro, Schrödinger, LLC, New York, NY, 2021) was utilised. Initially, a volume overlap matrix was generated to quantify the spatial similarity between ligand conformations, and the average linkage method was applied to this matrix. The optimal number of clusters was determined, and the ligand poses were grouped accordingly. A representative pose from the largest cluster was selected for subsequent analysis. The largest cluster, containing approximately 140 poses, was considered to represent the most plausible binding geometry of CBD within CYP9K1.

#### 2.10.3. MM-GBSA

Docking poses obtained by consensus docking analysis were subjected to binding free energy calculations performed using the Molecular mechanics/generalized Born surface area (MM-GBSA) method (Lyne et al., 2006) implemented in the Prime module of Maestro (Schrödinger Release 2021-2: Maestro, Schrödinger, LLC, New York, NY, 2021. Each calculation was performed using default settings. The OPLS4 force field was used for energy minimization, and the VSGB implicit solvation model was applied to estimate solvation effects.

#### 2.10.4. Induced fit docking

Induced Fit Docking (IFD) calculations were performed using the Induced Fit Docking protocol in Maestro (Schrödinger Release 2021-2: Maestro, Schrödinger, LLC, New York, NY, 2021; (Sherman et al., 2006). A representative ligand pose from the largest cluster obtained through consensus docking was selected as the center for grid generation. The IFD procedure was carried out using the standard protocol and generated multiple poses, approximately 75% of which closely matched the representative consensus docking structure.

### 2.11. Toxicity to non-target organisms

#### 2.11.1. Toxicity to bees (*Apis mellifera* Linnaeus)

##### 2.11.1.1 Collection of bees and test cages

Young adult worker honeybees from healthy, queen-right colonies with a known history and physiological status were collected from an apiary before the treatment. These colonies had not been treated with any chemicals, such as antibiotics or anti-Varroa agents, for five months preceding the study. Before the experiment, the bees were transferred from the apiary to an experimental room.

The tests were based on the following guidelines, considering necessary modifications to accommodate the specific properties of compounds: OECD Guideline No. 214 for the testing of chemicals: Honeybees, Acute Contact Toxicity Test (OECD, 1998) and existing EU testing guidelines (European Food Safety Authority, 2013).

The bees were divided randomly into groups of 15 and placed in plexiglass cages (7.1 x 12 x 12 cm) (Lusebrink, 2018). A hole on the upper side of each cage provided free access to the diet. The test cages were easy to clean and well-ventilated.

##### 2.11.1.2. Acute contact toxicity

Hemp-derived samples tested for toxicity were diluted in ethanol solvent carrier while the four common pyrethroid insecticides (tau-fluvalinate, deltamethrin, lambda cyhalothrin and beta cyfluthrin) were diluted in acetone. The compound solutions were applied using a Gilson micropipette (Gilson Inc., Middleton, WI, USA). Before the initiation of the treatment, the bees were placed in groups of 15 in cages. Bees from each cage were anesthetised for 15 seconds with carbon dioxide and placed on a Petri dish for topical application. A volume of 2 μL of the test solution was applied to the dorsal part of the thorax of each bee. After application, the bees were returned to their respective cages. Controls were conducted simultaneously and were treated with 2 μL of ethanol or acetone accordingly.

Following treatment, the bees had continuous access to a 50% (*w/v*) sucrose in water solution. Throughout the experiment, the honeybees were kept in an incubator, at a temperature of 25 ± 2°C and humidity maintained between 50-70%.

The bees were monitored for mortality every 24 hours post-treatment, for up to 96 hours.

#### 2.11.2. Toxicity on HaCaT and Caco-2 cell lines

##### 2.11.2.1. Cell culture

Human keratinocyte cell line (HaCaT) was cultured in Gibco DMEM medium, low glucose, supplemented with L -glutamine, penicillin-streptomycin and fetal bovine serum. Human intestinal epithelial cell line (Caco-2) was cultured in EMEM medium (supplemented with L –glutamine, non-essential amino acids, sodium pyruvate and fetal bovine serum. Cultures were maintained at 37 °C with a 5% CO_2_ humidified atmosphere.

##### 2.11.2.2. MTT assay

To evaluate cell viability, a methyl tetrazolium (MTT) assay was performed. Live cell count was determined using Trypan Blue staining and live cells were quantified using a Bürker chamber under a microscope. Caco-2 and HaCaT cells were cultured overnight in respective growth media at a 10,000 cells/well density in a 96-well plate. Cytotoxicity was determined after exposure of the cultures to dilutions 0–0.025% (*w/v*) of extract. Each extract was diluted in ethanol, the concentration of which did not exceed 0.05% in the final solution. The respective cell-culture medium was used as a control. After 24 hours of incubation, the medium was discarded, and MTT reagent was added to each well. Plates were further incubated at 37°C for 3 hours. Following the incubation period, DMSO was used to dissolve the formazan crystals. The color intensity was measured at 550 nm using a spectrophotometer (Halo LED 96, Dynamica Scientific, Livingston, UK). Fluorescence values lower than those of control cells indicated a reduction in the rate of cell proliferation. Conversely, a higher fluorescence rate indicated an increase in cell viability and proliferation. A biological replicate with 8 technical replicates per concentration and 16 technical replicates for the untreated control were included. Statistical analysis was performed using one-way ANOVA, with significance set at p < 0.05 and a confidence level of 95%.

## 3. Results

### 3.1. Cannabidiol inhibits CYP6CM1 and CYP9K1

For the initial screening, we selected thirty-seven extracts/fractions from twenty-one plant species from our in-house natural products library (Supplementary Table S1). Selection of extracts for testing was based on insecticidal properties described in the literature and the structural diversity of their major compounds. The library consisted mainly of crude methanolic (MeOH) and ethyl acetate (EtOAc) extracts, selected to broadly cover both polar/semi-polar metabolites (enriched in MeOH extracts) and more lipophilic constituents (enriched in EtOAc extracts). In addition, a limited number of enriched fractions were included and are detailed in Supplementary Table S1. We utilised *B. tabaci* CYP6CM1 (a known broad spectrum insecticide metaboliser) (Karunker et al., 2009; Nauen et al., 2013) for the primary screen due to the high yield we consistently achieve for this protein in heterologous expression, and verified positive results on our target enzyme, the *An. gambiae* CYP9K1.

In parallel, to prioritise potential inhibitors of *An. gambiae* CYP9K1, the in-house molecular library PHARMALAB (Fantel et al., 2020) consisting of approximately 1,870 small molecules was subjected to the following *in silico* workflow. The HHpred server was employed to identify the closest structurally characterized homolog of CYP9K1, Human Cytochrome P450 3A4. Virtual screening efforts were redirected toward compounds predicted to inhibit CYP3A4, under the hypothesis that scaffolds active against this homolog might also bind to CYP9K1. PHARMALAB was evaluated using three independent ADME prediction tools and 369 compounds were selected based on positive inhibition results against CYP3A4 across all three platforms. To further refine the selection while maintaining chemical diversity, the compounds were clustered by similarity and a representative molecule from each cluster was selected, yielding a final set of 39 candidates for further evaluation (Supplementary Table S3). Notably, CBD was included among the shortlisted candidates (Supplementary Table S3) and was therefore prioritised for experimental validation alongside the lead fraction identified in the extract-based screening.

In the *in vitro* extract/fraction screening, twenty of the thirty-seven samples tested contained compounds that interfered with the fluorescence detection in the inhibition assay precluding accurate measurements. For the remaining seventeen extracts moderate to low inhibition was observed (Supplementary Table S1). Notably, ethyl acetate extracts in most cases were more potent compared to methanolic extracts. This may reflect enrichment of more lipophilic constituents in the ethyl acetate extracts, consistent with the preference of P450 enzymes for lipophilic ligands (Feyereisen, 2005).

The most significant inhibition (32.32 ± 0.59%) was observed with the decarboxylated acidic fraction of a *C. sativa* SFE extract (SFEAFD) from the hemp variety ‘*Futura 75’*. To evaluate the influence of extraction method on inhibitory activity, we also prepared an extract from the same biomass using ethanolic UAE (ultrasound-assisted extraction), another standard and widely used method for cannabinoid enrichment (Citti et al., 2020). The corresponding ethanolic hemp extract was processed identically (Section 2.2), generating acidic, neutral, and decarboxylated acidic fractions, which were then tested alongside the SFE fractions (Table 1. List of screened ‘Futura 75’ hemp extracts and fractions, their respective percentage inhibition of Bemisia tabaci CYP6CM1 activity, and standard deviation (SD) at a concentration of 8.33 μg/mL (n=3).. The ethanolic extract and its corresponding fractions exhibited weaker inhibition against CYP6CM1, compared to the SFE extract and its fractions.

**Table 1.** List of screened ‘Futura 75’ hemp extracts and fractions, their respective percentage inhibition of Bemisia tabaci CYP6CM1 activity, and standard deviation (SD) at a concentration of 8.33 μg/mL (n=3).

| Tested hemp extracts/fractions | Inhibition<br>(% mean ± SD) |
| --- | --- |
| SFE total extract | $0.28 \pm 1.33$ |
| SFE acidic fraction | $12.58 \pm 3.00$ |
| SFE acidic fraction decarboxylated | $32.32 \pm 0.59$ |
| SFE neutral fraction | nd |
| EtOH total extract | nd |
| EtOH extract acidic fraction | 18.11 ± 5.72 |
| EtOH extract acidic fraction decarboxylated | 24.76 ± 3.06 |
| EtOH extract neutral fraction | 20.59 ± 3.56 |

*SFE: supercritical fluid extraction with CO<sub>2</sub>, EtOH: ethanolic, nd: not detected. Differences between SFE- and EtOH-derived neutral fractions are expected due to method-dependent chemical composition of the resulting fractions; thus, inhibition was detected for the EtOH neutral fraction but not for the SFE neutral fraction under these assay conditions.*

To identify the major constituents of the hemp SFEAFD fraction, which showed the highest CYP6CM1 inhibition, we performed UPLC-PDA and UHPLC-HRMS/MS analyses. UPLC-PDA identified CBD as the predominant component, constituting 27.68% of the fraction, with its acidic precursor CBDA present at 2.55% (Supplementary **Error! Reference source not found.**).

Since UPLC-PDA alone cannot definitively distinguish between certain neutral cannabinoids (Citti et al., 2018), the fraction was further analysed by UHPLC-HRMS/MS. Comparison with authentic reference standards confirmed CBD as the major cannabinoid and revealed five additional cannabinoids at trace levels (Figure 1 & Supplementary Table S4).

**Figure 1.**
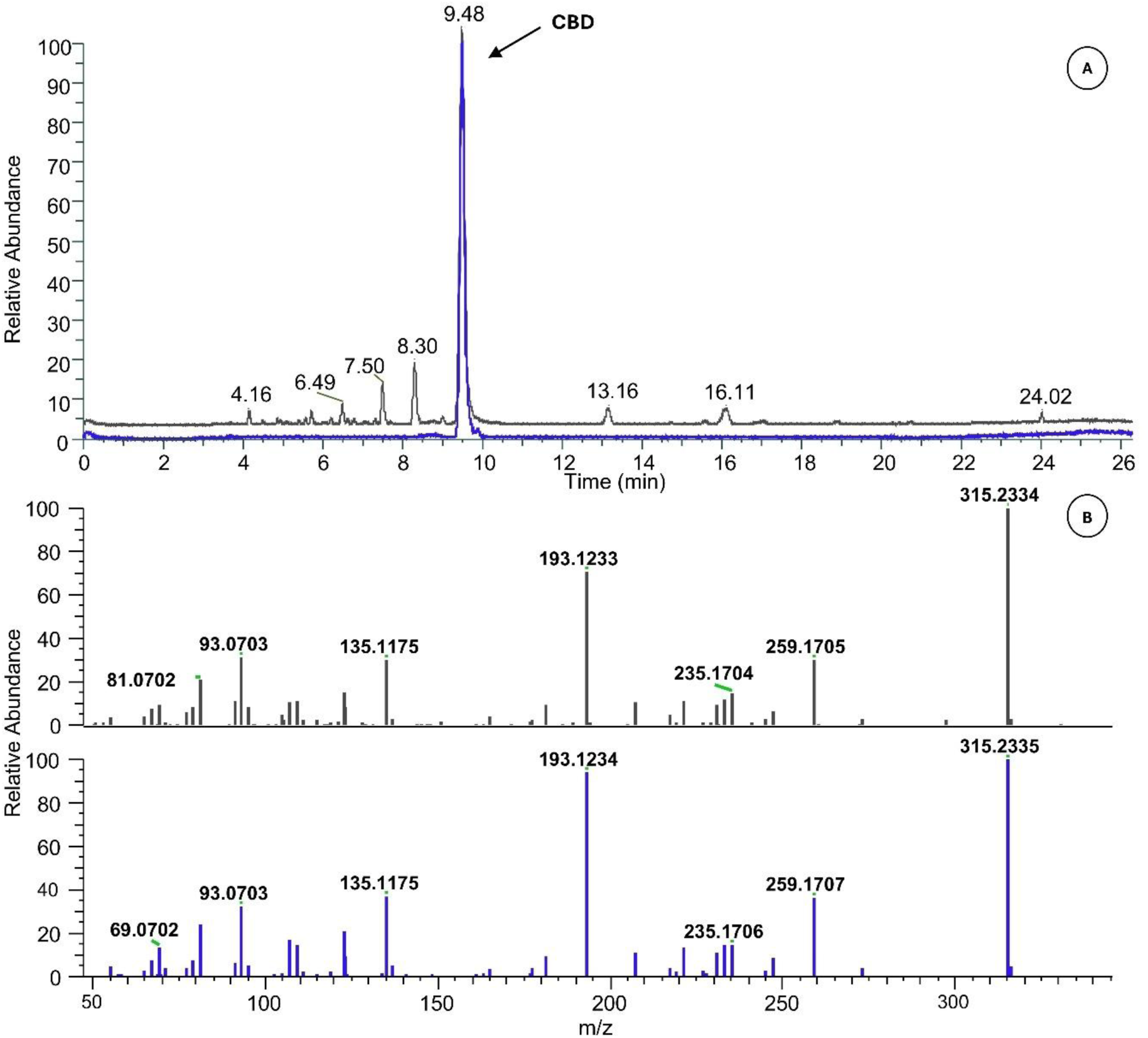
(A) Overlay of LC-HRMS base peak chromatograms of the decarboxylated acidic fraction of Cannabis sativa (black) and CBD reference standard (blue) in positive ionisation mode. CBD elutes at 9.48 min. (B) Stacked high resolution mass spectra of CBD from the decarboxylated acidic fraction (top) and the CBD reference standard (bottom) at 9.48 min.

Isolated CBD (99.9% purity) and its acidic form, CBDA, (99.6% purity) were tested *in vitro* for their inhibitory activity against CYP6CM1. CBD exhibited 64.92 ± 4.74% inhibition at 26 μM (approximately 8.2 μg/mL) and a -1.1°C shift to the enzyme’s *T*_m_ (melting temperature) in the differential scanning fluorimetry (DSF) assay performed (Supplementary Figure S2), while CBDA showed no inhibitory activity, indicating that decarboxylation is critical for inhibitory activity.

Next, the inhibitory properties of the decarboxylated acidic hemp fraction and CBD were tested against CYP9K1, a key insecticide detoxification enzyme in *An. gambiae*. At 8.33 μg/mL, the decarboxylated acidic hemp fraction exhibited mild inhibition of CYP9K1 (16.46 ± 5.24%) indicating the presence of bioactive constituents with weak enzyme-inhibitory properties. In contrast, pure CBD demonstrated substantially higher inhibition, reaching 52.79 ± 1.66% at 20 μM.

### 3.2. Semisynthetic CBD analogues and their CYP9K1 inhibition efficiency

To enhance the P450 inhibitory activity of CBD, a semisynthetic approach was employed. One of the two phenolic hydroxyl groups of CBD was selected as the main modification site. An O-alkyl ester intermediate was first prepared and used as a common precursor to introduce a set of heteroatom-rich side chains. Specifically, linear or branched aliphatic chains and heterocycle-containing substituents bearing a terminal amino group, with varying side-chain lengths, were installed at this position (Figure 2). These substituents were intended to modulate CBD’s polarity, conformational flexibility and to create additional opportunities for hydrogen bonds and electrostatic interactions within the P450 active site, while preserving the terpenoid core in order to explore structure–activity relationships (Bissantz et al., 2010).

**Figure 2.**
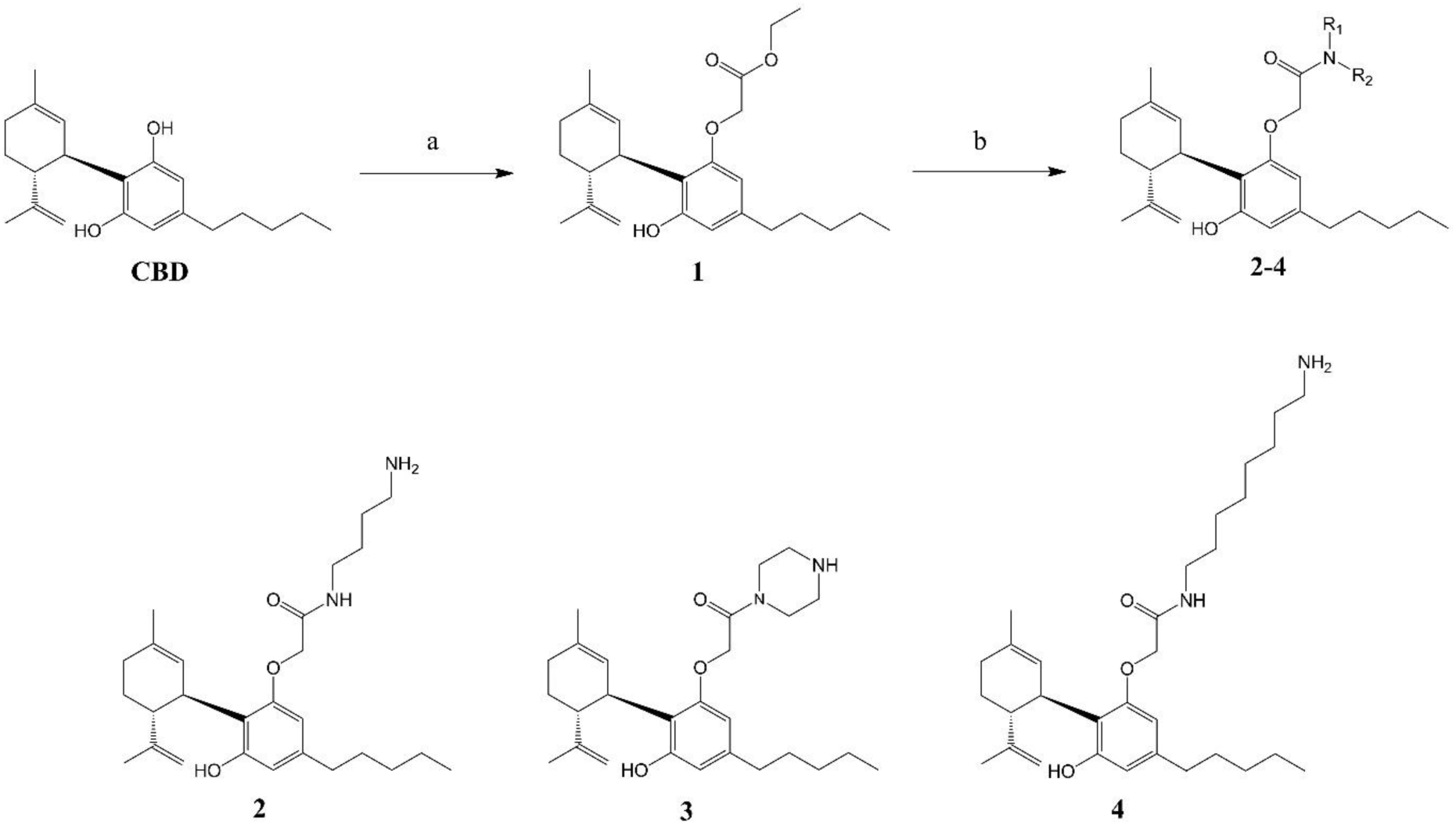
Chemical structures of CBD (parent scaffold) and its semisynthetic analogues. a. Ethyl bromoacetate, Cs_2_CO_3_, butanone, Ar, 90°C, 16h; b. 1,4-diaminobutane (for Analogue 2) or piperazine (for Analogue 3) or 1,8-diaminooctane (for Analogue 4), EtOH, Ar, 93°C, 16h.

The synthetic strategy for the preparation of the target compounds is shown in Figure 2. Briefly, CBD, obtained as described in Section 2.5 from *Cannabis sativa* L., was first subjected to O-alkylation with ethyl bromoacetate to afford the corresponding ester intermediate (Analogue 1). Subsequent nucleophilic substitution with the appropriate diamines provided the final CBD analogues 2–4.

When the semisynthetic analogues were screened at 20 μM, Analogue 3 exhibited the strongest effect, inhibiting CYP9K1 by 87.93 ± 2.16%, whereas the remaining analogues showed comparatively lower activities (all < 30% inhibition) (Figure 3 & Supplementary Table S5). CBD inhibited CYP9K1 by 52.79 ± 1.66% at the same concentration.

**Figure 3.**
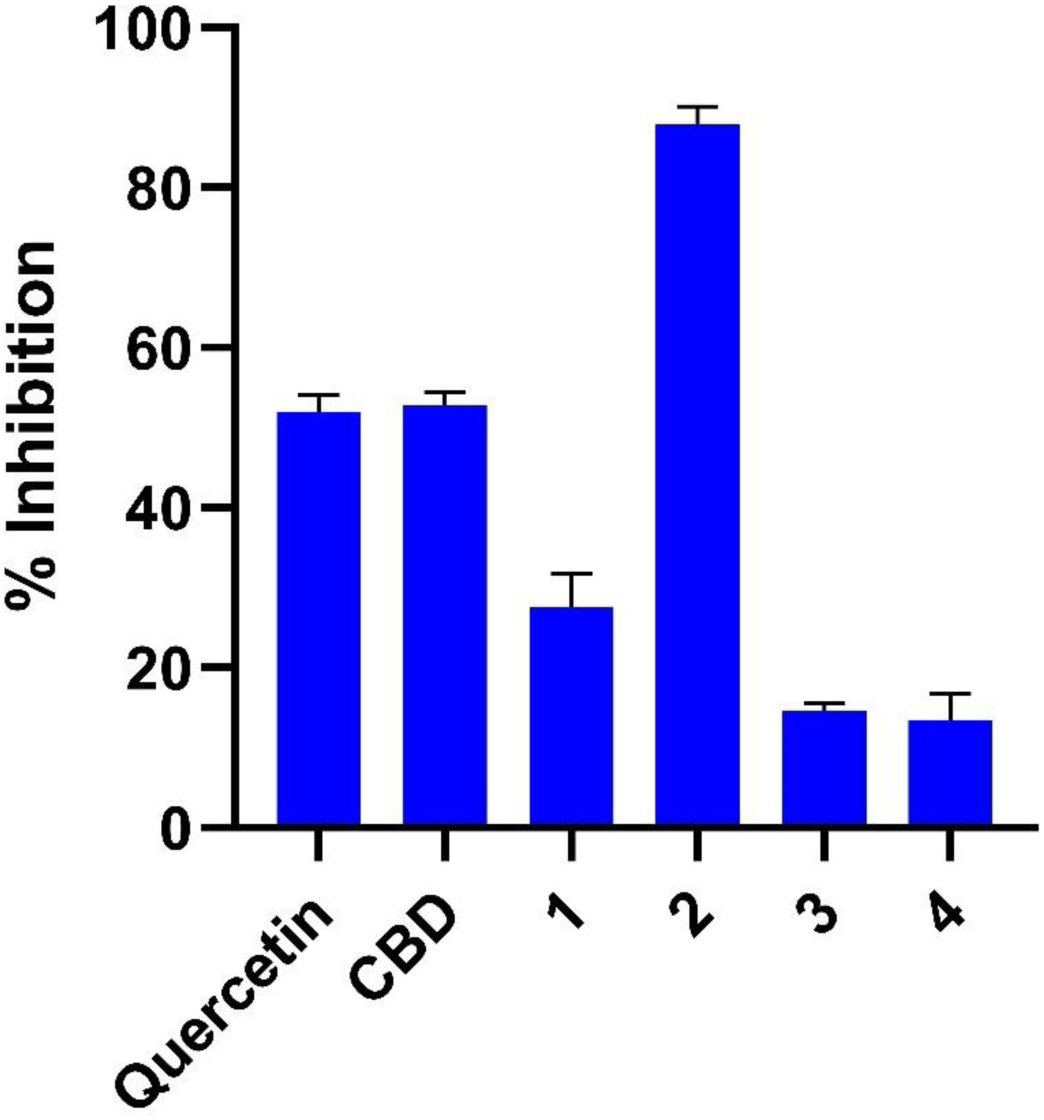
Bar graph comparing the percentage inhibition of CYP9K1 by CBD and its semisynthetic analogues, tested at 20 μM. Numbers indicate different CBD analogues. Quercetin was used as a positive control.

To quantify and compare inhibitory potency, concentration–response curves were generated for CBD and Analogue 3. The IC₅₀ for CBD was determined to be 18.37 μM, while CBD analogue 3 exhibited a significantly lower IC₅₀ of 2.50 μM (Figure 4).

**Figure 4.**
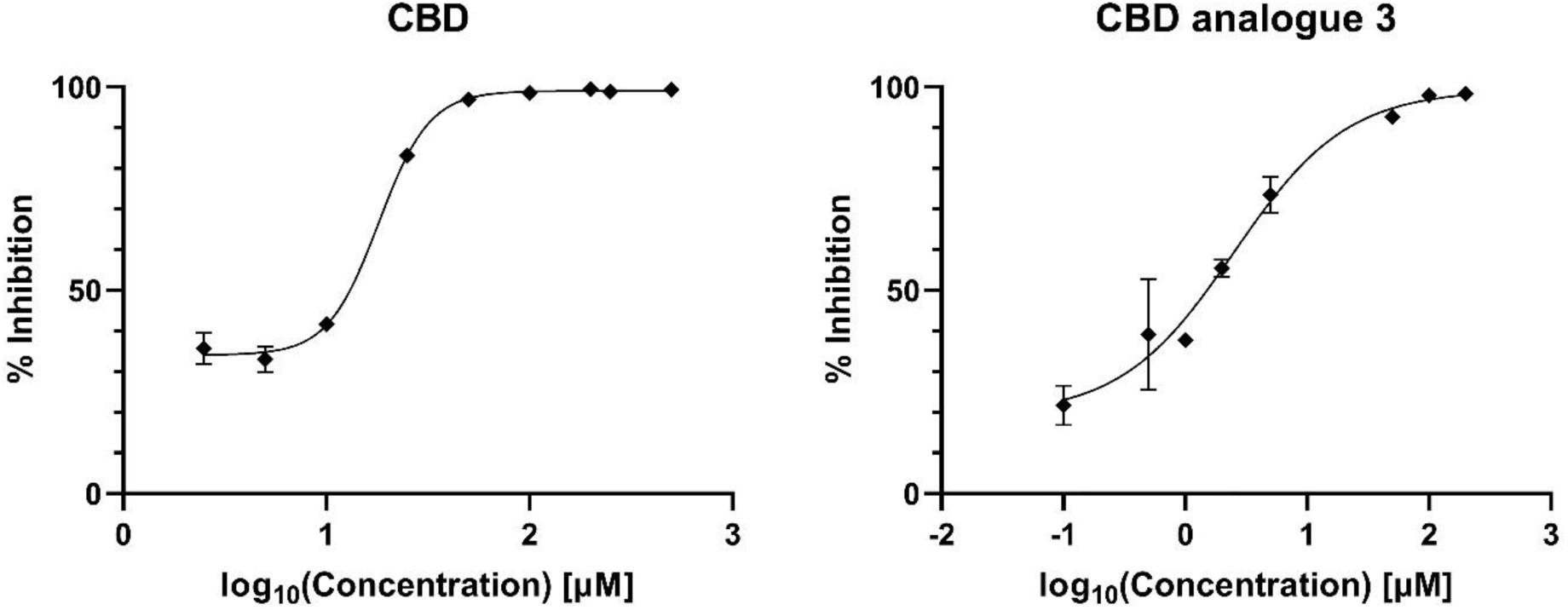
IC_50_ curves of CBD (left) and CBD analogue 3 (right) for inhibition of CYP9K1. Error bars represent standard deviation.

### 3.3. Evaluating the intrinsic toxicity of CBD and Analogue 3 in *An. gambiae* adults and their synergistic effect with deltamethrin

To further investigate the biological effects of the tested compounds, toxicity bioassays were conducted on *An. gambiae* adult mosquitoes. The first experiment assessed the intrinsic toxicity of the decarboxylated acidic hemp fraction, CBD and CBD analogue 3 in an insecticide-susceptible laboratory strain (G3). The second experiment evaluated their potential to synergise the toxic effects of deltamethrin in a pyrethroid-resistant *An. gambiae* colony (VK7).

None of the tested compounds exhibited marked intrinsic toxicity. Even at the highest concentration tested (20,000 ppm), mortality caused by CBD and Analogue 3 remained low (<30%) and was not significantly different from the solvent control (Supplementary). In contrast, as expected, deltamethrin caused substantial mortality (77% at 0.4 ppm). These findings indicate that CBD and its semisynthetic Analogue 3 do not possess strong direct insecticidal properties under the conditions tested.

We next evaluated whether CBD and Analogue 3 could enhance the toxic effect of deltamethrin. Mosquitoes from the VK7 strain, a pyrethroid-resistant colony, were treated topically with deltamethrin alone and in combination with CBD or Analogue 3 and mortality was assessed 24 h post-treatment.

At 3 ppm deltamethrin, the mean mortality in the VK7 strain was 16%, (consistent with its resistant phenotype). Co-application with 4,000 ppm CBD (the highest concentration tested that conferred ≤10% mortality in the toxicity test) resulted in a modest increase in mortality (35%); however, this difference did not reach statistical significance (Figure 5A). In contrast, co-application with 5,050 ppm (the highest concentration tested that conferred ≤10% mortality in the toxicity test) of Analogue 3 produced a statistically significant increase in mortality, from a mean of 23% when exposed to deltamethrin only to 65% when exposed to the combinatorial mixture (Figure 5B).

**Figure 5.**
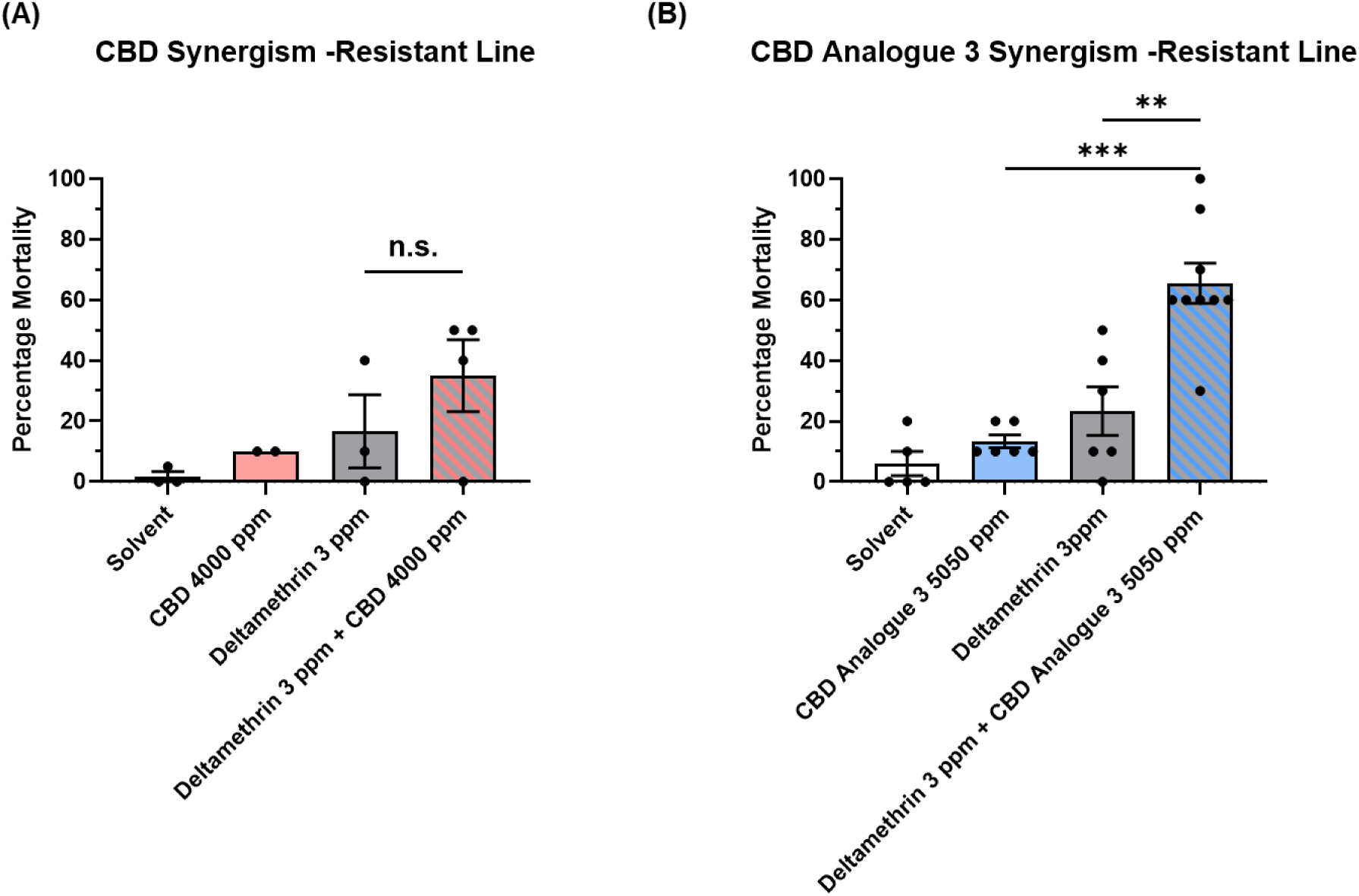
Percentage mortality of pyrethroid-resistant An. gambiae VK7 adults exposed to deltamethrin alone or in combination with CBD (A) or CBD analogue 3 (B). The synergistic effect was evaluated at concentrations of 3 ppm deltamethrin combined with 4,000 ppm CBD and 5,050 ppm CBD analogue 3. Solvent is a 1:1 mixture of ethanol and acetone. Error bars represent standard deviation.

### 3.4. *In silico* prediction of cannabidiol and Analogue 3 binding to CYP9K1

To investigate the molecular basis for CYP9K1 inhibition by CBD and to rationalise the enhanced potency of the most active semisynthetic derivative (Analogue 3), computational docking studies were performed. Cytochrome P450 enzymes are known for their highly flexible active sites and their ability to undergo significant conformational changes to accommodate a wide variety of substrates (Ekroos and Sjögren, 2006). The dynamic properties of P450 together with the absence of a CYP9K1 crystal structure, introduces uncertainty in accurately predicting ligand binding modes using conventional docking methods. To address these limitations, we set up a workflow primarily based on a consensus docking approach combined with IFD and binding free energy prediction (ΔG_bind_) as described in detail in the Methods section.

In brief, CBD was docked in two CYP9K1 AlphaFold-based models, generating 128 poses. Together with their energy-minimized structures, a total of 256 poses were selected for consensus analysis. The structures were clustered and representative structures of the most populated clusters were subjected to ΔG_bind_ prediction. AlphaFold models were generally more energetically favorable than those from the MODELLER structure and they were considered for further refinement. The representative pose of the largest cluster, containing approximately 140 poses, was subjected to Induced Fit Docking (IFD) calculations. ΔG_bind_ values were predicted for all IFD poses and the lowest energy structure was finally selected as the most plausible binding mode of CBD within the P450 cavity (Figure 6A). This orientation places 5-carbon aliphatic side chain directly above the heme iron with the first carbon of the chain adjacent to the ring located directly above the Fe²⁺ ion, a geometry that is critical for substrate oxidation. As an additional validation step of this theoretical model, it is noteworthy that Biotransformer 3.0 online tool predicts that human CYP3A4 oxidizes CBD specifically at this carbon (Supplementary Table S6; Djoumbou-Feunang et al., 2019).

**Figure 6.**
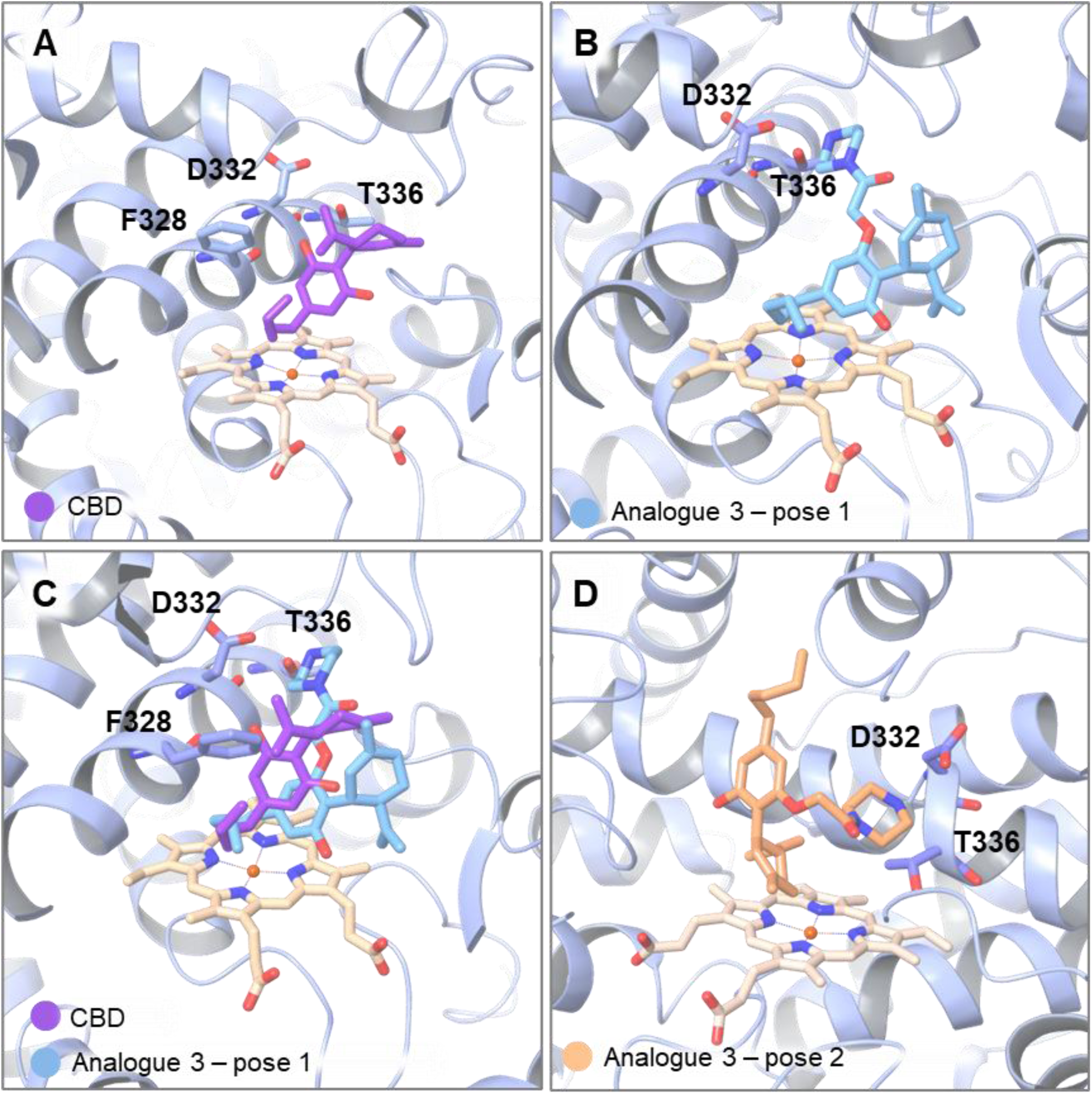
Docking poses of CBD and Analogue 3 in the predicted CYP9K1 model. (A) CBD illustrated in magenta is docked directly above the heme group. (B) Analogue 3 in light blue binds with its aliphatic benzene side chain positioned above the heme group, while the piperazine moiety is oriented near residues D332 and T336. (C) Overlay of CBD (magenta) and Analogue 3 (light blue), showing a similar binding geometry. (D) Most stable binding pose of Analogue 3, based on MM-GBSA ΔG_bind_ values.

We further investigated the binding of the most potent semisynthetic analogue, Analogue 3. For this purpose, IFD and MM-GBSA calculations were repeated using the same protocol. Among the resulting orientations, two poses were selected. One exhibiting a binding mode similar to that of CBD and another corresponding to the lowest predicted ΔG_bind_. In both orientations, the piperazine moiety is located similarly near residues D332 and T336, which might contribute to the higher affinity of Analogue 3 against CYP9K1 (Figure 6B, C, D). In the first pose (Figure 6B, C) the 5-carbon aliphatic chain is again oriented above the heme group and MM-GBSA prediction of ΔG_bind_ indicates that Analogue 3 binds more favorably than CBD, with a Δ_Gbind_ difference of approximately 4.6 kcal/mol, consistent with the observed in vitro potency. The most stable CYP9K1–Analogue 3 complex, illustrated in Figure 6D, positions the cyclohexene ring in close proximity to the heme group. Notably, the carbon atom located meta to the ring linker and ortho to the isopropenyl group lies near to the Fe²⁺ ion. This orientation is particularly compelling, as Biotransformer 3.0 also identifies this carbon as a likely site of oxidation by human CYP3A4. Although these models are based on predicted structures, they suggest that the enhanced activity of Analogue 3 may arise from its ability to form additional stabilizing interactions of the piperazine moiety within the CYP9K1 active site.

### 3.5. Testing the toxicity of compounds to non-target organisms

Toxicity of the decarboxylated acidic hemp fraction (SFEAFD), CBD and Analogue 3 to adult bees was evaluated by applying the compounds topically to their thorax and estimating the dose that causes 50% mortality (LD_50_) at multiple post-exposure time points (Supplementary tables S5A-D). At the final assessment, four days after treatment, the extract and the two cannabis compounds showed significantly lower toxicity compared to the pyrethroids deltamethrin and beta-cyfluthrin and comparable (for the analogue) or lower (for the fraction and CBD) levels of toxicity to the pyrethroids Lambda-cyhalothrin and tau-fluvalinate.

To test potential toxicity of CBD and the SFE decarboxylated acidic fraction to humans toxicity assays were conducted on HaCaT (human keratinocytes) and Caco-2 (human intestinal epithelial) cell lines

In HaCaT cells, both the SFE acidic fraction and CBD did not show any effect on cell’s viability when applied at a concentration of 0.001% *w/v* (10 ppm). However, significant toxicity was observed when the concentration was doubled (0.002% *w/v*; 20 ppm) (Figure 7A). At this and higher concentrations, a considerable reduction in cell viability was observed, with the fraction reducing viability by 92% and pure CBD by 96%.

**Figure 7.**
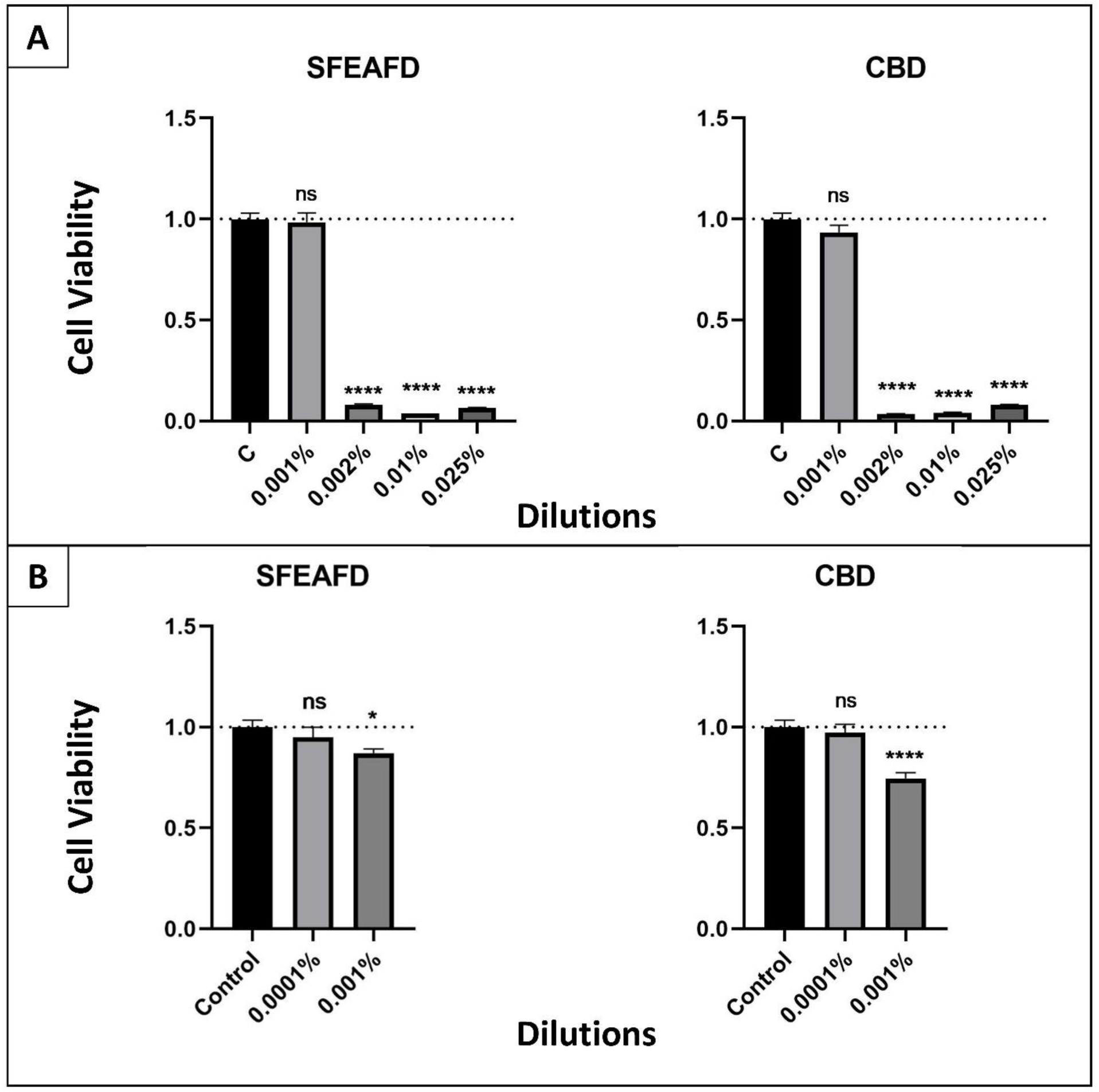
Cytotoxicity of the decarboxylated acidic hemp fraction (SFEAFD) and pure CBD in human cell lines. (A) HaCaT cell viability after 24 h exposure (MTT assay). Cells were treated with SFEAFD or CBD at 0.025%, 0.01%, 0.002%, and 0.001% (w/v). Significant cytotoxicity (****p < 0.0001) was observed at 0.025%, 0.01%, and 0.002%; no significant effect (ns) was detected at 0.001%. (B) Caco-2 cell viability after 24 h exposure (MTT assay). Cells were treated with SFEAFD or CBD at 0.001% and 0.0001% (w/v). A significant reduction in viability occurred at 0.001% for both SFEAFD (*p < 0.05) and CBD (****p < 0.0001); no effect was seen at 0.0001%. Data are presented as mean ± standard deviation (SD).

Cytotoxicity was also evaluated in Caco-2 cells to assess potential toxicity upon oral exposure. At the lowest tested concentration (0.0001% *w/v* or 1 ppm), no cytotoxic effects were observed for either the fraction or pure CBD, indicating that at trace levels, these compounds are unlikely to pose a significant risk to intestinal epithelial cells (Figure 7B). However, at 0.001% *w/v*, a mild but statistically significant reduction in cell viability was detected. Specifically, the fraction exhibited 13% cytotoxicity, whereas CBD showed a more pronounced effect, reducing cell viability by 25%.

## 4. Discussion

Over-expression of P450s that bind and detoxify insecticides is one of the best characterised metabolic resistance mechanisms in mosquitoes and other insects. Blocking the activity of those enzymes can substantially restore susceptibility and increase the efficacy of insecticidal formulations. The use of PBO, a P450 inhibitor, has been widely adopted in malaria control interventions, where it is being co-applied with pyrethroid insecticides on bed-nets and is considered a major resistance management strategy (Gleave et al., 2021). To preserve our ability to manage metabolic resistance, it is important to have an arsenal of synergists that either perform better than PBO or can be used in alternation or as backup in cases of emergence of resistance.

Plant extracts and their constituents have proven a valuable source not only of insecticidal compounds, but also of compounds that can inhibit metabolic enzymes and act as synergists. For example ethanol senescent leaf extracts of *Jatropha gossypifolia* L. and *Melia azedarach* L. not only deter *Spodoptera frugiperda* Giles larvae, but also enhance cypermethrin toxicity, partly through inhibition of P450s and esterases (Bullangpoti et al., 2011). Combinations of *Solanum xanthocarpum* Solanum Schrad. & Wendl. extracts with cypermethrin similarly increase larvicidal activity against *Culex quinquefasciatus* Say, indicating synergistic interactions (Mohan et al., 2006). Aqueous extracts of *Piper amalago* L. var. *amalago* and *Kalanchoe pinnata* (Lam.) Pers. inhibit key *An. gambiae* P450s (CYP6P9a, CYP6M2, CYP6P3) enhancing permethrin activity (Francis et al., 2025). More mechanistically resolved examples include *Rhinacanthus nasutus* (L.) Kurz naphthoquinones (rhinacanthins), which inhibit pyrethroid-metabolising CYP6AA3 and CYP6P7 and synergise cypermethrin toxicity in *S. frugiperda* cells expressing these enzymes (Pethuan et al., 2012), as well as *Andrographis paniculata* (Burm.f.) Wall. ex Nees leaf extracts and isolated flavones that act as potent CYP6AA3/CYP6P7 inhibitors and increase cypermethrin susceptibility *in vitro* (Kotewong et al., 2014).

To date, only one study has examined CBD’s effects on insect detoxification enzymes, showing that dietary CBD reduces food consumption, growth and overall CYP450 activity in *S. frugiperda* larvae (Abendroth et al., 2023). Our study provides a comprehensive analysis of CBD and a semisynthetic CBD analogue as plant-derived scaffolds for the development of P450 inhibitors synergising the action of neurotoxic insecticides.

The initial screening against recombinant CYP6CM1 showed that a decarboxylated acidic fraction of a *C. sativa* (‘Futura 75’) SFE-CO_2_ extract was the most active in our library. Subsequent UPLC-PDA and UHPLC-HRMS/MS analyses established that this fraction is enriched in neutral cannabinoids, with CBD as the predominant constituent.

Our screening indicated that the supercritical fluid extract (SFE) of *C. sativa* ‘Futura 75’ yielded a more potent inhibitory fraction than an ethanolic ultrasound-assisted extract (UAE) of the same biomass (Table 1). This likely reflects the higher selectivity of supercritical CO₂ for less polar, lipophilic compounds, such as cannabinoids (Moreno et al., 2020), which are known to interact with cytochrome P450 enzymes (Nasrin et al., 2021; Sevrioukova, 2025).

Cannabis exhibits a highly complex phytochemistry, with over 545 identified secondary metabolites, including cannabinoids, terpenoids, flavonoids, sterols and lipids (Jin et al., 2020). Hemp SFE extracts contain highly lipophilic compounds, including various lipid constituents (Martinez et al., 2023). Liquid-liquid extraction (LLE) further proved to be an effective method for separating lipids from cannabinoids by manipulating the pH, allowing neutral, non-ionisable compounds (e.g. lipids and waxes) to remain into the organic phase (neutral fraction), while enriching the aqueous phase (acidic fraction) with naturally abundant acidic cannabinoids (e.g. CBDA). LLE has been widely applied for cannabinoid enrichment in analytical and preparative contexts (Lu et al., 2023). Herein, part of the acidic fraction was decarboxylated to produce the corresponding neutral cannabinoids, which were also assessed for P450 inhibitory activity. Notably, cannabinoids are biosynthesised and predominantly occur in plants and fresh extracts as carboxylic acids (e.g. CBDA), which are less characterised and generally exhibit lower biological potency. Decarboxylation converts these into neutral forms (e.g., CBD), which are more extensively studied and biologically active (Izzo et al., 2009; Mechoulam and Hanuš, 2000).

Accordingly, in our study, the decarboxylated acidic fraction of the SFE extract, enriched in neutral cannabinoids, particularly CBD, displayed the strongest inhibition of CYP6CM1. When tested as pure compounds, CBD caused substantial inhibition of CYP6CM1 at micromolar concentrations (approximately 65% inhibition at 26 μM), while its acidic precursor CBDA showed negligible activity, indicating that the neutral cannabinoids are primarily responsible for the observed P450 interactions. CBD also inhibited *An. gambiae* CYP9K1 with comparable potency, whereas the crude hemp fraction showed only mild activity.

Together, these results identify CBD as the principal P450-active component of the SFE-derived hemp fraction (SFEAFD) and demonstrate that it can bind to and inhibit P450s from different insect species. In parallel, the results also underscore the advantage of combining selective extraction (e.g., SFE-CO₂) with pH-controlled fractionation and subsequent decarboxylation to enrich bioactive neutral cannabinoids from crude hemp extracts.

The inhibition efficiency of CBD (18.37 μM) is within the range reported for other plant-derived P450 inhibitors, although some isolated natural compounds have been reported to act at lower micromolar concentrations. For instance, purified naphthoquinones (rhinacanthins) from *Rhinacanthus nasutus* inhibit related lepidopteran P450s at approximately 10 µM (Pethuan et al., 2012).

CBD is well documented to bind to various mammalian cytochrome P450 enzymes, where it commonly acts as a competitive or mixed inhibitor while also serving as a low-efficiency substrate for certain isoforms. This dual behavior is characteristic of ligands that occupy positions near the heme iron and retain sufficient residence time to interfere with substrate turnover (Bansal et al., 2022; Doohan et al., 2021; Nasrin et al., 2021). CBD binding is primarily mediated by hydrophobic interactions of the terpenoid scaffold within the active site, with oxidisable moieties positioned close to the catalytic center, and is often accompanied by relatively weak or transient polar contacts with surrounding residues (Kundu et al., 2025; Sevrioukova, 2025). Consistent with this behavior, CBD has been shown to inhibit several mammalian CYPs in the low micromolar range, suggesting that its scaffold provides productive active-site access but leaves scope for affinity enhancement through targeted functionalisation.

Based on these general features of CBD–CYP interactions, a semisynthetic strategy was employed to enhance P450 inhibitory activity by introducing substituents capable of strengthening enzyme–ligand interactions. One of the two phenolic hydroxyl groups of CBD was selected as the primary site for modification, as derivatisation at this position is known to be well tolerated and allows installation of additional functionality without disrupting the terpenoid core required for active-site engagement (Kinaci et al., 2025). An O-alkyl ester intermediate was prepared as a common precursor to enable the introduction of a small series of heteroatom-rich side chains. Specifically, linear or branched aliphatic chains and heterocycle-containing substituents bearing a terminal amino group, with varying side-chain lengths, were incorporated. These modifications were designed to modulate molecular polarity and conformational flexibility, and to promote additional hydrogen-bonding and electrostatic interactions within the P450 active site while preserving the terpenoid core in order to explore structure–activity relationships (Bissantz et al., 2010).

Among the semisynthetic compounds, Analogue 3, bearing a piperazinyl moiety, emerged as the most potent inhibitor of CYP9K1, with an IC₅₀ of 2.50 μM compared with 18.37 μM for CBD, an approximately seven-fold gain in potency. This magnitude of improvement is consistent with successful optimisation campaigns for other natural product-derived P450 inhibitors. For instance, a monocarbonyl curcumin derivative exhibited a three-fold lower IC₅₀ than curcumin against *B. tabaci* CYP6CM1 (Ioannou et al., 2025). Similar strategies, guided by inhibition assays against conserved enzyme targets like CYP3A4, have led to significant potency gains for other plant-derived synergist scaffolds, such as dillapiole (Francis Carballo-Arce et al., 2019).

The *in silico* analysis provides a plausible structural rationale for the observed inhibitory activities of CBD and its optimised analogue. Docking studies with CYP9K1 models indicate that CBD binds with its pentyl side chain oriented toward the catalytic heme iron, positioning a benzylic carbon—predicted as a potential oxidation site for human CYP3A4—in proximity to the Fe²⁺ ion. For the more potent CBD analogue 3, the most stable binding pose suggests a reoriented geometry that places the cyclohexene ring near the heme while the introduced piperazine moiety engages in stabilising interactions with residues D332 and T336. These predicted orientations are consistent with the significantly more favorable calculated binding free energy for Analogue 3 compared to CBD, corroborating its lower experimental IC₅₀ value.

Despite the limitations imposed by using homology-based CYP9K1 models and the known flexibility of P450 active sites, the docking results offer working hypotheses on how the orientation of CBD’s pentyl side chain and the introduced piperazinyl substituent of Analogue 3 may contribute to the observed inhibition efficiencies.

We also tested CBD and Analogue 3 for their toxic and synergistic effect in adult *An. gambiae* mosquitoes, as this life stage is the primary target of insecticide-based interventions. None of the compounds was intrinsically toxic, given that less than 30% mortality was observed at the 20,000 ppm dose. CBD did not display statistically significant synergism with deltamethrin in the pyrethroid-resistant *An. gambiae* VK7 strain (that has P450 based resistance) (Williams et al., 2019), despite a modest trend towards increased mortality. In contrast, the CBD-derived analogue 3 substantially enhanced deltamethrin-induced mortality at 5,050 ppm (approximately 1.01 µg/mosquito), by approximately 2.8fold.

When the outcomes are expressed as a co-toxicity factor (Norris et al., 2018) to enable comparison with other studies using this metric, we obtain a value of 80. This value is higher than the reported co-toxicity factors for PBO in pyrethroid-resistant *Ae. aegypti* (25 with permethrin in Baker et al., 2023 and 22 with deltamethrin in Norris et al., 2018), even though PBO was applied topically at higher doses (2 µg/mosquito and 10,000 ppm, respectively) than Analogue 3. The co-toxicity factor of 80 is also higher than that reported for clove leaf essential oil (68), which exhibited the highest value among all essential oils tested in resistant *Ae. aegypti* and is lower than the co-toxicity factor reported for the monoterpenoid fenchone (160) when co-applied with permethrin at 2 µg/mosquito. Additional structural modifications of analogue 3 could further enhance its potency.

To obtain an initial assessment of the safety profile of CBD and Analogue 3 toward non-target organisms, we conducted contact toxicity tests on adult honeybees. Both compounds were less toxic to adult worker bees than several pyrethroids used in agriculture and public health and lower or equally toxic to tau-fluvalinate, a pyrethroid currently used in beekeeping applications for controlling Varroa mites. These results indicate that CBD and Analogue 3 do not pose a greater immediate hazard to honeybees than established synthetic pyrethroids.

The toxicity of CBD was also evaluated in human cell lines. Cell viability of HaCaT cells was not affected at a concentration of 0.001% *w/v*. In contrast, Caco-2 cells were more sensitive, exhibiting a 25% reduction in viability at the same concentration. These results align with previous studies indicating that CBD, while bioactive, may exhibit toxic effects at relatively high concentrations (Gingrich et al., 2023; Huestis et al., 2019). Notably, while CBD toxicity data is emerging, similar cytotoxicity profiles for commonly used insecticide synergists such as PBO remain largely unreported in human skin and intestinal cell lines. This gap in the literature highlights the need for comparative toxicological assessments of not only the active compounds but also of their adjuvants. Furthermore, the clear concentration-dependent cytotoxicity observed at higher doses indicates that translation to field applications would require careful toxicological evaluation.

Although *C. sativa* is extensively referenced in the literature for its insecticidal properties (Ona et al., 2022), this is the first time it is used as a scaffold for yielding insecticide synergists. In addition, most previous work on *C. sativa* has focused on hemp essential oils. Essential oil studies demonstrate that various hemp chemotypes can kill larvae of *Aedes*, *Anopheles* and *Culex* species (Abé et al., 2018; Bedini et al., 2016; Benelli et al., 2018; Mazzara et al., 2023; Pavela, 2009; Rossi et al., 2020; Wanas et al., 2020), but volatility and short residual activity limit their direct use as long-term control agents. In contrast, the current work employed cannabinoid-enriched supercritical fluid extracts (SFE) and ethanolic extracts from *C. sativa* inflorescences (var. ‘Futura 75’), rich in stable, lipophilic compounds with greater longevity. Furthermore, the synthesis of CBD derivatives is straightforward and economical, since we have established a robust method of isolating CBD, and the subsequent semisynthesis is relatively simple.

Overall, our findings expand the chemical space for P450-mediated insecticide resistance management and highlight CBD analogues as promising candidates with a favorable, in many aspects, profile for enhancing the efficacy of pyrethroids in the control of disease-vector insects.

## CRediT authorship contribution statement

**Maria Chalkiadaki:** Data Curation, Formal analysis, Investigation, Methodology, Visualization, Writing - Original Draft **Linda Grigoraki:** Data Curation, Formal analysis, Methodology, Writing - Review & Editing **Dimitra Tsakireli:** Investigation, Methodology, Writing - Review & Editing **Georgia Vasalaki:** Investigation, Visualization **Petros S. Tzimas:** Formal analysis, Investigation, Methodology, Visualization, Writing - Review & Editing **Mengling Chen:** Investigation, Visualization **Latifa Remadi:** Investigation **Rino Ragno:** Formal analysis, Methodology, Software, Writing - Review & Editing **Ifigeneia Akrani:** Investigation, Software, Writing - Review & Editing **Emmanouel Mikros:** Data Curation, Supervision, Validation, Writing - Review & Editing **Rafaela Panteleri:** Formal analysis, Investigation, Methodology, Visualization **Spyros Vlogiannitis:** Formal analysis, Investigation, Methodology **Vassilios Myrianthopoulos:** Data Curation, Validation, Visualization, Writing - Review & Editing **Ioannis K. Kostakis:** Methodology, Supervision, Validation, Writing - Review & Editing **Leandros A. Skaltsounis:** Data Curation, Resources, Supervision, Writing - Review & Editing **John Vontas:** Conceptualization, Funding acquisition, Project administration, Resources, Supervision, Validation, Writing - Review & Editing **Maria Halabalaki:** Conceptualization, Funding acquisition, Project administration, Resources, Supervision, Validation, Writing - Review & Editing

All authors have read and agreed to the published version of the manuscript.

## Declaration of Competing Interest

The authors declare that they have no known competing financial interests or personal relationships that could have appeared to influence the work reported in this paper.

## Acknowledgements

This publication is based on research funded by: the Hellenic Foundation for Research and Innovation (H.F.R.I.) under the H.F.R.I. call “Basic research Financing (Horizontal support of all Sciences)” under the National Recovery and Resilience Plan “Greece 2.0” funded by the European Union – NextGenerationEU (H.F.R.I. Project Number:016044) and the European Union’s MSCA-RISE-2020 program CypTox- 101007917.

## Appendix A. Supplementary material

## Data Availability

Data will be made available on request.

## Notes

### Competing Interest Statement

The authors have declared no competing interest.

