## Supplementary material for "Mitigating CYP450-Mediated Insecticide Resistance in Malaria Vectors with Cannabis-Derived Synergists: The Potential of Cannabidiol"

Table S1. Inhibitory activity of screened plant extracts and fractions against Bemisia tabaci CYP6CM1. Data include botanical family, species, plant part, and extraction method/fraction type. Activity is reported as mean percentage inhibition ± standard deviation (SD) at 8.33 µg/mL (n = 3).

| **Species** | **Family** | **Plant part** | **Extract/ fraction type** | **Inhibition**  **(% mean ± SD)** |
| --- | --- | --- | --- | --- |
| *Alkanna graeca* | Boraginaceae | whole plant | MeOH (UAE) | 2.35 ± 1.14 |
| *Artemisia arborescens* | Asteraceae | aerial parts | EtOAc (UAE) | nd |
|  |  |  | THF (UAE) | nd |
| *Cannabis sativa* | Cannabaceae | inflorescences | SFE CO_2-_LLE acidic fraction | 12.58 ± 3.00 |
|  |  |  | SFE CO_2-_LLE acidic fraction decarboxylated | 32.32 ± 0.59 |
|  |  |  | SFE CO_2-_LLE neutral fraction | nd |
| *Centaurea attica* | Asteraceae | whole plant | EtOAc (UAE) | nd |
|  |  |  | MeOH (UAE) | nd |
| *Centaurea peucedanifolia* | Asteraceae | flowering aerial parts | EtOAc (UAE) | 17.10 ± 5.10 |
| *Daphne laureotica* | Thymelaeaceae | branches and leaves | EtOAc (UAE) | 12.39 ± 2.10 |
|  |  |  | MeOH (UAE) | 3.40 ± 0.76 |
| *Daphne oleoides* | Thymelaeaceae | branches and leaves | EtOAc (UAE) | nd |
|  |  |  | MeOH (UAE) | nd |
| *Daphne sericea* | Thymelaeaceae | branches and leaves | EtOAc (UAE) | 20.87 ± 0.99 |
|  |  |  | MeOH (UAE) | 6.47 ± 1.20 |
| *Dianthus haematocalyx* ssp. *pindicola* | Caryophyllaceae | whole plant | EtOAc (UAE) | 0.37 ± 0.30 |
|  |  |  | MeOH (UAE) | nd |
| *Epilobium hirsutum* | Onagraceae | aerial parts | EtOAc (UAE) | nd |
|  |  |  | MeOH (UAE) | nd |
| *Gentiana lutea* | Gentianaceae | roots | MeOH (UAE) | nd |
| *Morus alba* | Moraceae | wood | MeOH (UAE) | 29.94 ± 9.72 |
| *Onobrychis alba* ssp. *laconica* | Fabaceae | perennial aerial parts and roots | EtOAc (UAE) | 21.33 ± 0.23 |
|  |  |  | MeOH (UAE) | 7.45 ± 1.76 |
| *Onobrychis alba* ssp. *laconica* | Fabaceae | whole plant | EtOAc (UAE) | 28.36 ± 0.06 |
|  |  |  | MeOH (UAE) | 10.72 ± 0.23 |
| *Pistacia lentiscus* | Anacardiaceae | resin | acidic triterpenic fraction after LLE | nd |
|  |  | leaves | MeOH fraction after XAD7 | nd |
| *Rubia peregrina* | Rubiaceae | aerial parts | EtOAc (UAE) | nd |
|  |  |  | MeOH (UAE) | nd |
| *Salvia candidissima* | Lamiaceae | whole plant | EtOAc (UAE) | 16.11 ± 0.53 |
|  |  |  | MeOH (UAE) | 6.67 ± 1.00 |
| *Sesamum indicum* | Pedaliaceae | seed | oil LLE- ACN fraction | nd |
| *Thapsia garganica* | Apiaceae | aerial parts | EtOAc (UAE) | nd |
|  |  |  | MeOH (UAE) | nd |
| *Olea europea* | Oleaceae | fruits | Total polyphenolic fraction from EVOO after LLE | 25.68 ± 0.01 |
| *Verbascum arcturus* | Scrophulariaceae | annual aerial parts | EtOAc (UAE) | nd |
|  |  |  | MeOH (UAE) | nd |

*UAE: ultrasound-assisted extraction, MeOH: methanolic, EtOAc: ethyl acetate, THF: tetrahydrofuranic, SFE: supercritical fluid extraction, LLE: liquid-liquid extraction EtOH: ethanolic, ACN: acetonitrile, EVOO: extra virgin olive oil, nd: not determined.*

Table S2. Root-mean-square deviations (RMSDs) of the CYP3A4 experimental binding sites with those of the CYP9K1 models from Alpha Fold (AF) and MODELLER (modbase).

| **PDB Entry code** | **AF** | **modbase** |
| --- | --- | --- |
| 7KVK | 3.1 | 5.1 |
| 9COY | 3.0 | 4.8 |
| 6UNL | 3.0 | 4.4 |
| 6UNJ | 3.3 | 5.7 |
| 6UNI | 2.9 | 5.2 |
| 6UNM | 4.2 | 8.9 |
| 6BDH | 3.5 | 8.8 |
| 7KVM | 3.0 | 4.5 |
| 6BD7 | 3.5 | 5.0 |
| 7KVI | 3.1 | 4.5 |

Table S3. 39 representative compounds from in-silico screening using SwissADME, ADMET Predictor (Simulation Plus Inc.), and ADMETlab platforms.

| 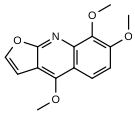 | 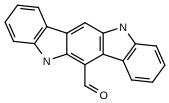 | 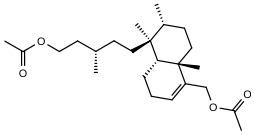 |
| --- | --- | --- |
| 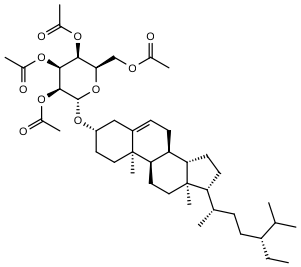 | 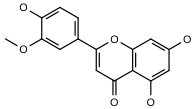 | 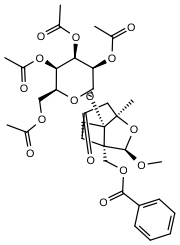 |
| 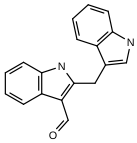 | 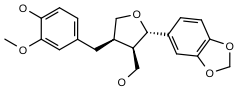 | 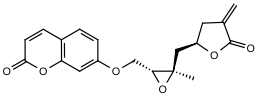 |
| 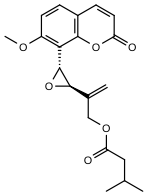 | 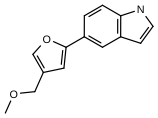 | 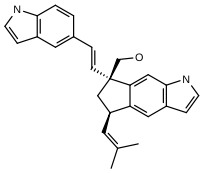 |
| 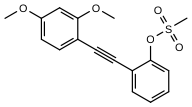 | 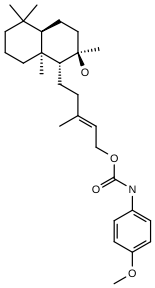 | 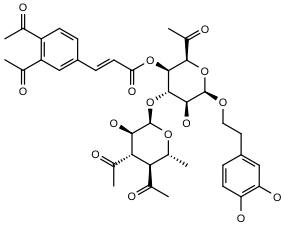 |
| 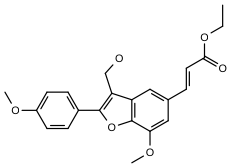 | 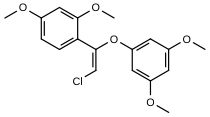 | 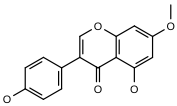 |
| 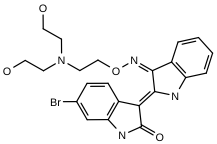 | 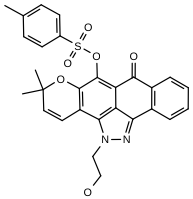 | 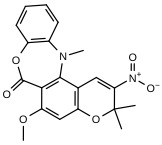 |
| 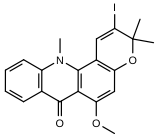 | 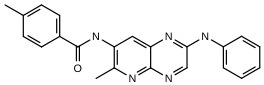 | 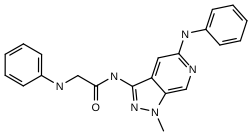 |
| 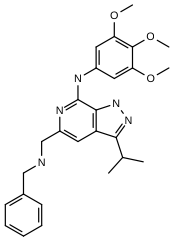 | 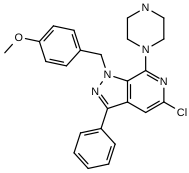 | 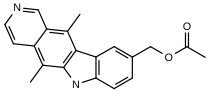 |
| 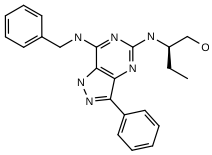 | 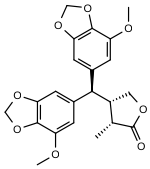 | 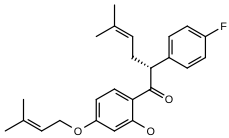 |
| 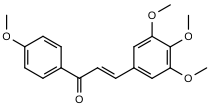 | 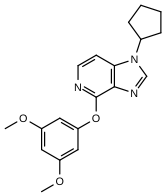 | 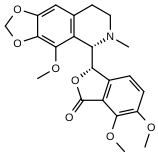 |
| 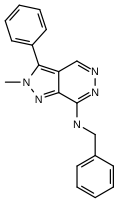 | 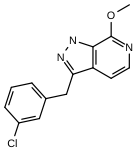 | 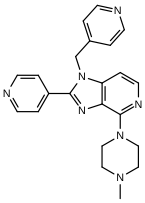 |
| 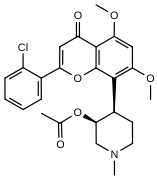 | 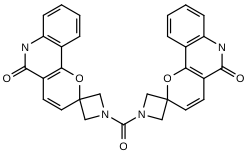 | 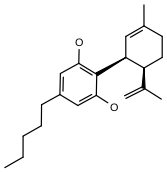 |

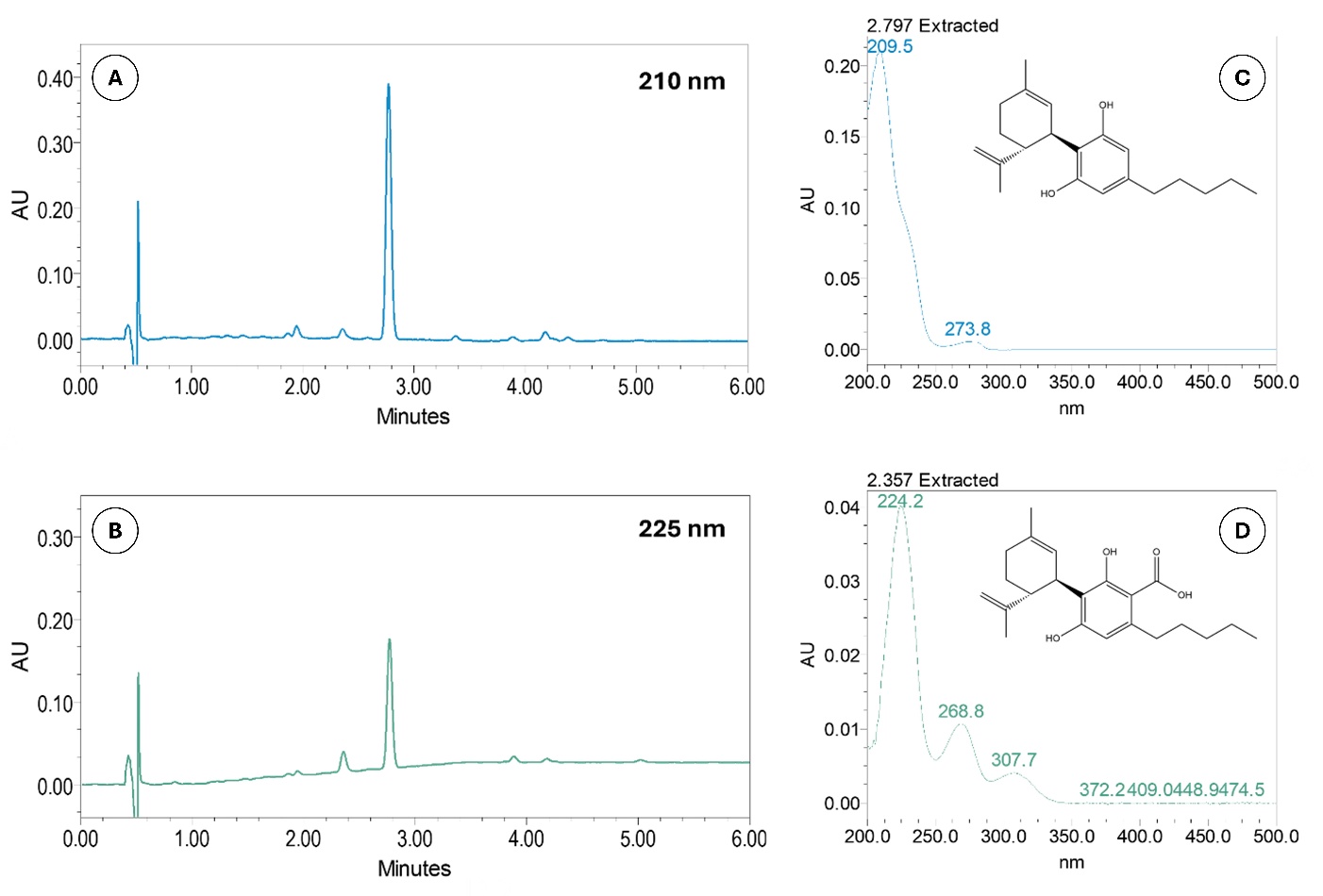

Figure S1. UPLC-PDA chromatograms of the decarboxylated acidic hemp fraction at 0.10 mg/mL, recorded at (A) 210 nm and (B) 225 nm. UV/Vis absorbance spectra of (C) CBD and (D) CBDA.

Table S4. UHPLC-HRMS/MS data of the decarboxylated acidic hemp fraction (SFEAFD) annotated cannabinoids in comparison to reference standards.

| **Compound** | **CBDV** | **CBE** | **CBDA** | **CBD** | **CBN** | **THC** | **CBC** | **CBCT** |
| --- | --- | --- | --- | --- | --- | --- | --- | --- |
| **Rt (min)** | 6.51 | 7.50 | 8.30 | 9.48 | 13.17 | 16.09 | 20.72 | 24.02 |
| **Adduct** | [M+H]^+^ | [M+H]^+^ | [M+H]^+^ | [M+H]^+^ | [M+H]^+^ | [M+H]^+^ | [M+H]^+^ | [M+H]^+^ |
| **Experimental *m/z*** | 287.2005 | 331.2262 | 359.2212 | 315.2314 | 311.2000 | 315.2312 | 315.2316 | 315.2317 |
| **Elemental composition (EC)** | C_19_H_26_O_2_ | C_21_H_30_O_3_ | C_22_H_30_O_4_ | C_21_H_30_O_2_ | C_21_H_26_O_2_ | C_21_H_30_O_2_ | C_21_H_30_O_2_ | C_21_H_30_O_2_ |
| **RDB_eq_. values** | 6.5 | 6.5 | 7.5 | 6.5 | 8.5 | 6.5 | 6.5 | 6.5 |
| **Δm (ppm)** | -0.20 | -1.64 | -1.29 | -1.31 | -1.85 | -2.08 | -0.71 | -0.63 |
| **HRMS/MS fragments** | 165.0909 (100) 287.2004 (97) 231.1378 (34) 135.1167 (31) 93.0698 (30) 81.0698 (22) 95.0491 (19) 91.0542 (17) 207.1379 (16) | 205.1222 (100) 109.1011 (68) 135.0440 (37) 331.2267 (17) | 341.2110 (100) 219.1016 (36) 261.1483 (14) | 315.2317 (100) 193.1223 (82) 135.1167 (35) 93.0698 (34) 259.1692 (33) 123.0439 (22) 81.0698 (22) 95.0490 (19) 235.1691 (16) (107.0855 (16) 233.1535 (15) | 223.1128 (100) 311.2020 (92) 293.1913 (41) 241.1235 (30) 195.1178 (20) 178.0785 (16) | 315.2317 (100) 193.1223 (87) 259.1691 (363) 93.0698 (35) 135.1168 (34)  81.0698 (25) 123.0440 (19) 95.0491 (18) 91.0542 (18) | 193.1223 (100) 81.0698 (45) 259.1693 (32) 315.2319 (26) 69.0699 (21) 233.1537 (18) 93.0699 (17) 135.1168 (16) 109.1011 (16) 123.0441 (15) | 315.2318 (100) 193.1223 (39) 259.1692 (19) 135.1167 (18) |

Rt: retention time expressed in minutes; RDBeq: ring and double bond equivalent; Δm: mass accuracy expressed in ppm.

To further assess the possibility of direct CBD binding to CYP6CM1, the biophysical method of differential scanning fluorimetry (DSF) was employed (Niesen et al., 2007). The effect of ligand binding on the enzyme thermal stability was monitored in terms of the change in melting temperature (*T*_m_) and results revealed that CBD induced a -1.1°C shift to the enzyme’s *T*_m_ compared to the apoenzyme, suggesting a weak yet measurable binding interaction between CYP6CM1 and CBD (Supplementary Figure S2).

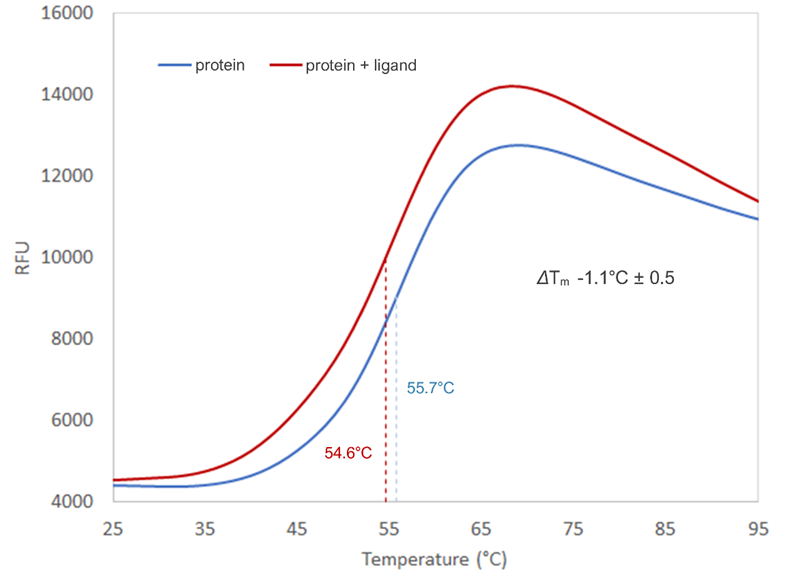

Figure S2. DSF curve showing the unfolding status of the target protein (CYP6CM1) in the absence (blue) and presence (red) of a ligand (CBD). The difference in the melting temperature is indicated as ΔT_m_.

Table S5. List of screened CBD analogues, their respective percentage inhibition of Anopheles gambiae CYP9K1 activity, and standard deviation (SD) at 20 μΜ (n=3).

| **Compound** | **IUPAC names** | **Inhibition**  **(% mean** ± **SD)** |
| --- | --- | --- |
| CBD | 2-[(1*R*,6*R*)-3-methyl-6-prop-1-en-2-ylcyclohex-2-en-1-yl]-5-pentylbenzene-1,3-diol | 52.79 ± 1.66 |
| Analogue 1 | ethyl 2-(((1'R,2'R)-6-hydroxy-5'-methyl-4-pentyl-2'-(prop-1-en-2-yl)-1',2',3',4'-tetrahydro-[1,1'-biphenyl]-2-yl)oxy)acetate | 27.50 ± 4.26 |
| Analogue 2 | N-(4-aminobutyl)-2-(((1'R,2'R)-6-hydroxy-5'-methyl-4-pentyl-2'-(prop-1-en-2-yl)-1',2',3',4'-tetrahydro-[1,1'-biphenyl]-2-yl)oxy)acetamide | 14.62 ± 0.91 |
| Analogue 3 | 2-(((1'R,2'R)-6-hydroxy-5'-methyl-4-pentyl-2'-(prop-1-en-2-yl)-1',2',3',4'-tetrahydro-[1,1'-biphenyl]-2-yl)oxy)-1-(piperazin-1-yl)ethanone | 87.93 ± 2.16 |
| Analogue 4 | N-(8-aminooctyl)-2-(((1'R,2'R)-6-hydroxy-5'-methyl-4-pentyl-2'-(prop-1-en-2-yl)-1',2',3',4'-tetrahydro-[1,1'-biphenyl]-2-yl)oxy)acetamide | 7.01 ± 5.73 |

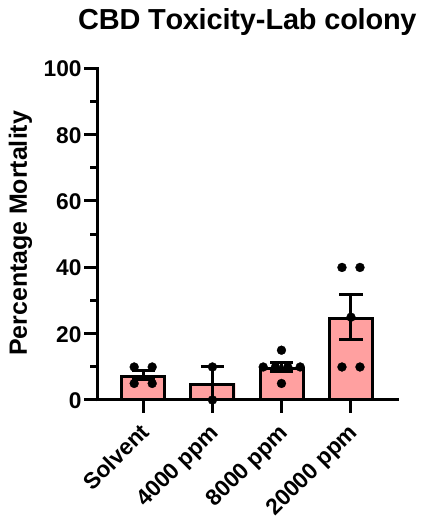

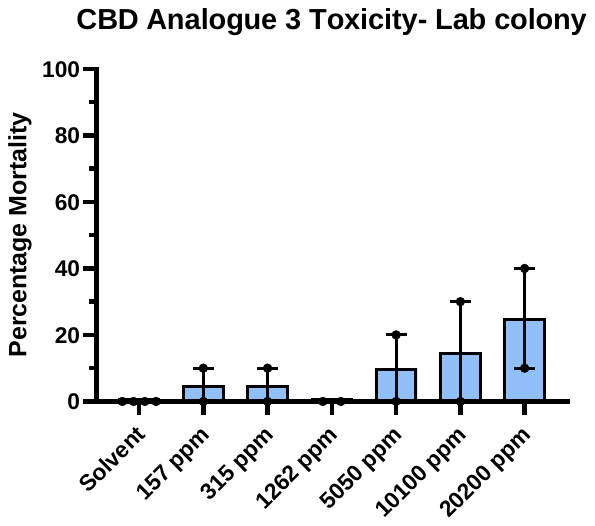

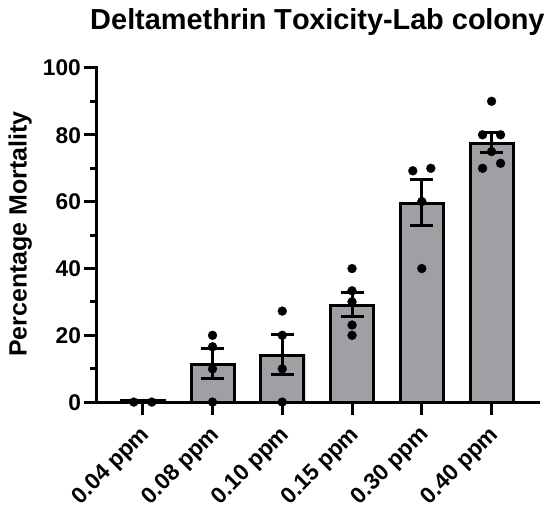

Figure S3. Percentage mortality of insecticide-susceptible Anopheles gambiae adults exposed to increasing concentrations of CBD, CBD analogue 3, and deltamethrin. Error bars represent standard deviation.

Table S6. Human P450 3A4 Oxidation sites of CBD and analogue 3 as predicted by online platform BioTransforemer 3.0.

| Compound | Structure | CYP3A4 BioTransformer 3.0 Oxidation Site |
| --- | --- | --- |
| CBD | 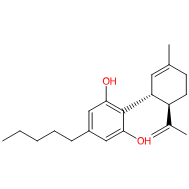 | 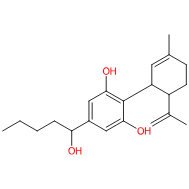 |
| Analogue_3 | 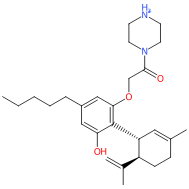 | 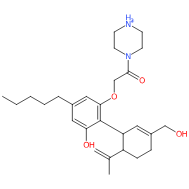 |

Table S7. Acute contact toxicity (LD₅₀, μg/bee) of test substances in Apis mellifera at 24 (A), 48 (B), 72 (C), and 96 h (D) post-treatment (95% CI; relative toxicity index).

| (A) 24 hours post treatment | | | |
| --- | --- | --- | --- |
| Test substance | LD_50_ μg/bee | 95% Confidence Interval (μg/bee) | Relative toxicity index |
| SFEAFD | **>10** | - | >16.38 |
| CBD | **>20** | - | >8.19 |
| Analogue 3 | **>20** | - | >8.19 |
| Tau-Fluvalinate | **163.89** | 99.99 to 258.88 | - |
| Deltamethrin | **2.39** | 1.35 to 4.09 | 68.57 |
| Lambda-cyhalothrin | **15.20** | 8.66 to 27.64 | 10.78 |
| Beta-cyfluthrin | **0.50** | 0.331 to 0.745 | 327.78 |
| (B) 48 hours post treatment | | | |
| Test substance | LD_50_ μg/bee | 95% Confidence Interval (μg/bee) | Relative toxicity index |
| SFEAFD | **>10** | - | >6.05 |
| CBD | **16.45** | 12.760 to 26.004 | 3.83 |
| Analogue 3 | **11.17** | 7.554 to 18.766 | 5.64 |
| Tau-Fluvalinate | **63.05** | 38.01 to 100.41 | - |
| Deltamethrin | **0.68** | 0.12 to 5.19 | 92.72 |
| Lambda-cyhalothrin | **8.43** | 5.00 to 14.66 | 7.48 |
| Beta-cyfluthrin | **0.20** | 0.135 to 0.289 | 315.25 |
| (C) 72 hours post treatment | | | |
| Test substance | LD_50_ μg/bee | 95% Confidence Interval (μg/bee) | Relative toxicity index |
| SFEAFD | **>10** | - | >1.4 |
| CBD | **10.20** | 8.291 to 12.673 | 1.41 |
| Analogue 3 | **5.77** | 2.983 to 7.803 | 2.49 |
| Tau-Fluvalinate | **14.39** | 7.53 to 25.30 | - |
| Deltamethrin | **0.18** | 0.11 to 0.28 | 79.95 |
| Lambda-cyhalothrin | **3.56** | 2.05 to 6.38 | 40.42 |
| Beta-cyfluthrin | **0.139** | 0.087 to 0.209 | 103.55 |
| (D) 96 hours post treatment | | | |
| Test substance | LD_50_ μg/bee | 95% Confidence Interval (μg/bee) | Relative toxicity index |
| SFEAFD | **>10** | - | - |
| CBD | **7.474** | 5.583 to 9.264 | > 1.33 |
| Analogue 3 | **3.297** | 1.162 to 4.718 | > 3.03 |
| Tau-Fluvalinate | **3.69** | 1.91 to 6.49 | > 2.71 |
| Deltamethrin | **0.044** | 0.022 to 0.070 | > 227.27 |
| Lambda-cyhalothrin | **1.42** | 0.85 to 2.46 | > 7.04 |
| Beta-cyfluthrin | **0.074** | 0.041 to 0.117 | > 135.13 |
